# ChatGEM: An Agentic Architecture Enabling Interactive Simulation of Genome-Scale Metabolic Models

**DOI:** 10.64898/2026.07.20.739662

**Authors:** Niaz Bahar Chowdhury, August George, Sumit Purohit, Angela Cintolesi, Erin Bredeweg, Jeffrey Czajka, Kelly Stratton, Yuqian Gao, Molly Stephenson, Joshua R. Elmore, Allan Scott, Damon Leach, Abby Jerger, Teresa Lemmon, Paul Piehowski, Kylee Tate, James M. Fulcher, Alex Beliaev, Kristin E. Burnum-Johnson, Paul Rigor, Jaydeep P. Bardhan

## Abstract

Genome-scale metabolic models (GEMs) are powerful tools for predicting cellular phenotypes and guiding microbial strain engineering, yet broad adoption remains challenging due to the computational expertise required. To overcome that, we present ChatGEM, an agentic platform that enables interactive GEM simulation through natural language. Built on the multi-agent ADEPT framework, ChatGEM integrates COBRApy within a retrieval-augmented generation (RAG) architecture that coordinates code generation and execution through specialized agents. Benchmarking across three tasks of increasing complexity showed that RAG-enabled code generation improved the mean overall performance score from 2.63 to 4.20 while reducing the execution time significantly starting from routine to complex tasks. Application of ChatGEM using an enzyme-constrained GEM (ecGEM) for four engineered *Pseudomonas putida* KT2440 strains identified the constitutive strain as the optimal chassis for succinate overproduction using a succinate leakage index - a prediction observed experimentally. Therefore, ChatGEM democratizes metabolic modeling by enabling researchers without computational expertise to perform sophisticated GEM-based analyses through natural language, and, hence, accelerating scientific discovery.

## INTRODUCTION

Genome-scale metabolic models (GEMs) have become foundational tools in systems biology, enabling the quantitative simulation of cellular metabolism under diverse genetic and environmental conditions (Kim, et al., 2017). Since their inception, GEMs have been applied to diverse applications including predicting growth phenotypes, identifying essential genes, guiding rational strain design, and interpreting multi-omics data in organisms ranging from *Escherichia coli* to human cell lines (Alsiyabi, et al., 2022; Heirendt, et al., 2019). The introduction of enzyme-constrained genome-scale metabolic models (ecGEMs), which incorporate proteomics data and enzyme kinetic parameters to enforce a finite total protein budget, substantially improved predictive accuracy over GEMs and enabled the study of proteome allocation under different conditions (Sanchez, et al., 2017; Shahreen, et al., 2024; Shahreen, et al., 2025). Moreover, computational strain design methods, most notably OptKnock (Burgard, et al., 2003), formalized the coupling of biomass formation with target chemical production, establishing growth-coupled overproduction as a central paradigm in metabolic engineering.

Despite these valuable developments, the application of GEM-based analysis remains concentrated among computational biologists. Executing a standard workflow, whether running flux balance analysis (FBA) (Orth, et al., 2010), formulating a strain design problem, or reconstructing an ecGEM, requires significant computational expertise in COnstraints-Based Reconstruction and Analysis (COBRA) framework (Ebrahim, et al., 2013), accessible through Python or Matlab, and familiarity with biochemical databases, such as KEGG (Kanehisa, et al., 2025) and ModelSEED (Seaver, et al., 2021). These requirements constitute a substantial entry barrier for many biologists, biochemists, and metabolic engineers, requiring significant computational training. The consequence is a persistent gap between the analytical potential of GEM-based methods and their actual adoption across the synthetic biology community.

The rapid development of large language models (LLMs) (Vaswani, et al., 2023) has opened a new avenue for bridging this gap. LLMs have demonstrated increasingly capable performance on scientific reasoning, code generation, and multi-step problem solving, and their deployment in agentic architectures, where autonomous agents coordinate tool use, memory, and iterative refinement, has enabled end-to-end automation of complex analytical workflows (Huang, et al., 2025). In biology, this has given rise to a wave of domain-specific tools. For instance, CRISPR-GPT automated gene-editing experimental design through an LLM agent integrating domain expertise and retrieval techniques (Qu, et al., 2026). Next, CellWhisperer established natural language as an interface for single-cell RNA-seq exploration through multimodal contrastive learning on over one million transcriptomes (Schaefer, et al., 2025). Moreover, BioAgents demonstrated that RAG-enhanced multi-agent systems built on small language models could match human expert performance on conceptual genomics tasks (Mehandru, et al., 2025). D2Cell applied LLMs to mine metabolic engineering strategies from over 10,000 literature entries, coupling the extracted knowledge to a hybrid deep learning and GEM-based target prediction model (Li, et al., 2026). Most recently, LLMs were directly used for GEM reconstruction workflows to streamline the curation of Human2, a consensus human GEM with tissue- and organ-specific resolution (Luo, et al., 2026).

However, to our knowledge, no existing tool has yet provided a conversational interface capable of handling the full stack of quantitative GEM simulations across the broad range of modeling tasks. Here we present ChatGEM, an LLM-based coding agent that enables interactive simulation and analysis of GEMs through natural language. Built on the ADEPT (George, et al., 2025) multi-agent framework, ChatGEM integrates COBRApy based tools within a retrieval-augmented generation (RAG) architecture (Lewis, et al., 2020) that coordinates code generation, execution, and interpretation through specialized agents. We benchmarked ChatGEM across three tasks of increasing biological complexity, from routine FBA with targeted reaction knockouts, to OptKnock strain design, to a fully integrated enzyme-constrained thermodynamic pathway optimization (ecOptMDFPathway), an ecGEM version of OptMDFPathway (Hadicke, et al., 2018), demonstrating that RAG-grounded code generation improved mean Overall Performance Score (OPS) from 2.63 to 4.20 across all three tasks. The agentic architecture substantially reduced time requirements across all tasks, with ChatGEM-generated code executing up to 65 times faster than manual coding workflows and eliminating the iterative debugging overhead that consumes significant expert time in complex modeling tasks. Through a biological case study using *P. putida* KT2440, ChatGEM reconstructed eight ecGEMs across four engineered strains and two time points, identified EDD, GLCDpp, and ACONTa as dominant protein sinks in the central carbon pathway, correctly predicting the Constitutive strain as the optimal chassis for succinate overproduction prior to experimental validation. All of these were accomplished through natural language interaction, without the user writing any code, demonstrating that ChatGEM can compress workflows that previously required specialized computational effort into a single conversational session.

## RESULTS

### ChatGEM Agentic Framework

ChatGEM is a specialized agentic system built from the ADEPT framework and tailored for GEM code generation and execution, integrating COBRApy, StrainDesign, and commercial solvers like Gurobi alongside ADEPT’s general scientific capabilities. ChatGEM inherits from ADEPT’s full architecture, which is designed for security, scalability, and flexibility (Figure 1).

**Figure 1:**
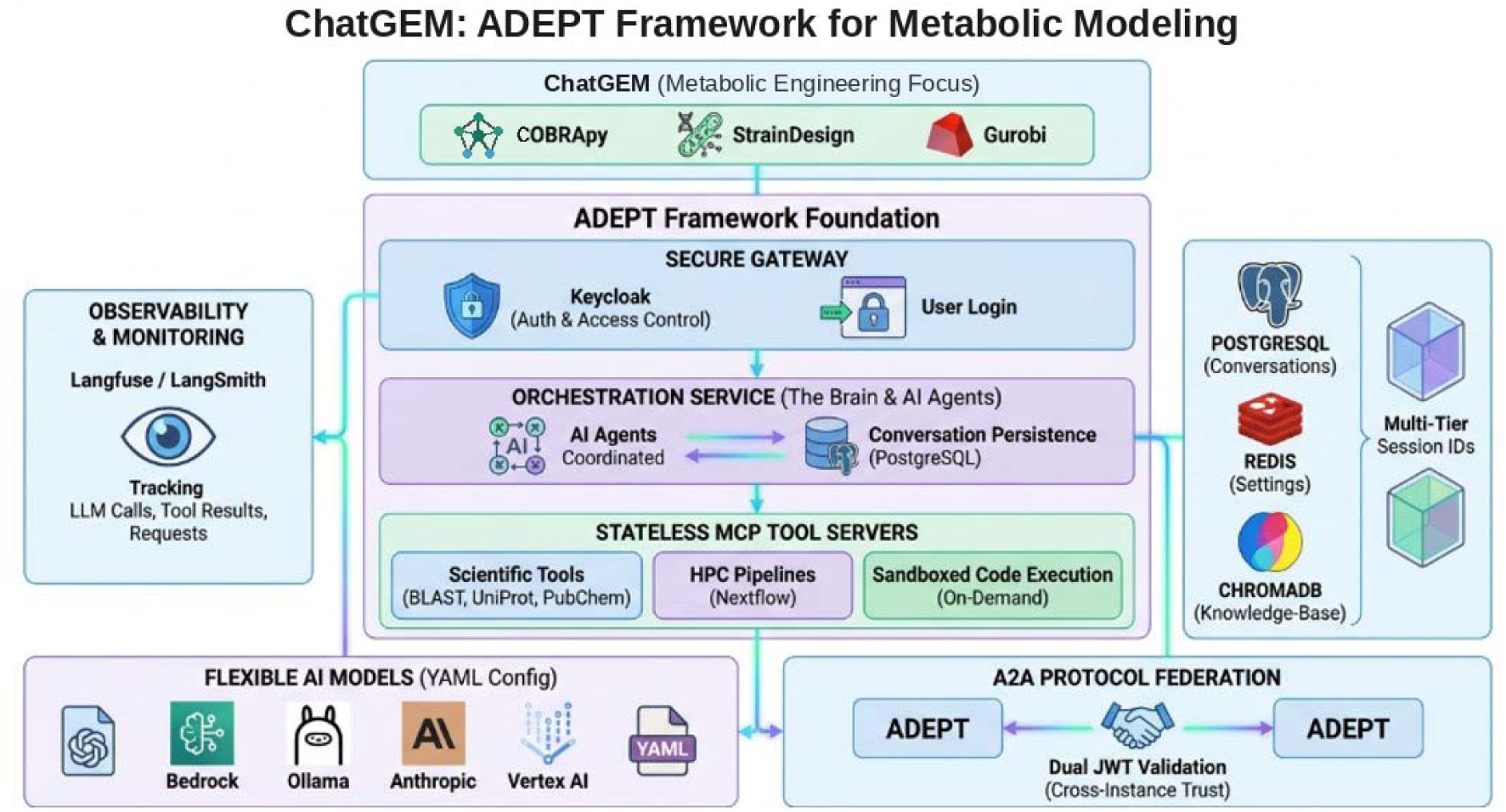
ChatGEM and ADEPT system design overview. ChatGEM adds common metabolic engineering tools (COBRApy, StrainDesign) to the ADEPT framework. ADEPT is LLM-provider agnostic and includes core components for secure access, orchestration, and MCP tools. Additionally, ADEPT supports observability/monitoring, external data stores, and agent interoperability.

ADEPT is built on a three-tier design: a secure gateway that handles user login and access control via Keycloak; a central orchestration service that acts as the coordinator for AI agents and persisting conversations in PostgreSQL; and a set of stateless MCP tool servers that run scientific tools (BLAST, UniProt, PubChem), HPC pipelines (Nextflow), and sandboxed code on demand.

Conversations, tool settings, and knowledgebase data are stored across specialized backends (PostgreSQL, Redis, and ChromaDB), with multi-tier session IDs keeping conversations, tool calls, and multi-agent workflows isolated. Retrieval augmented generation (RAG) is supported on demand via automated ingestion into the vector database. LLM calls and tool results can be tracked using Langfuse or LangSmith as well as a baseline Docker logging system. These tools support observability and monitoring of LLM requests, enabling improved auditability and debugging support.

ADEPT is designed to be model agnostic and with support for common model providers such as OpenAI, Anthropic, and Ollama via a YAML config file. For ChatGEM, the default option remains o4-mini for its smaller size and reasoning capability. Instances of ADEPT can also securely federate with one another via the A2A protocol using dual JWT validation for cross-instance trust. This enables secure agent federation across on-prem and cloud providers. Finally, the ADEPT uses containerization across the services to ensure a modular and scalable design.

### Efficient and Fast Systems Biology Code Generation and Execution

A central objective of ChatGEM is to generate and execute complex GEM workflows through natural language. To test this, we benchmarked ChatGEM across three tasks of varying computational complexity using the *P. putida* KT2440 GEM (iJN1463) (Nogales, et al., 2020): FBA with targeted reaction knockouts; strain design via OptKnock; and a combined enzyme- and thermodynamic-constrained modeling (ecOptMDFPathway). Each task was executed under two conditions, with and without RAG, to directly quantify the contribution of domain-specific context retrieval to code quality and biological correctness.

Generated code was evaluated by Claude Sonnet 4.6 with regards to two metrics (Supplementary File 1). Coding Accuracy (1–5) captured whether the code produced the correct biological outputs, including flux distributions, knockout assignments, target metabolite production rates, and best COBRApy and python coding practices. Code Completeness (1–5) assessed structural quality: proper use of library APIs, modular organization, interpretable variable naming, and sufficient documentation for independent reproduction. These scores were combined into an Overall Performance Score (OPS):

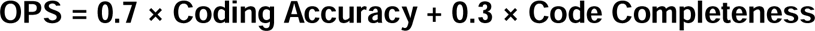

For the simplest task, FBA with reaction knockouts, both conditions produced working code, but important differences in correctness emerged. Without RAG, ChatGEM identified the succinate exchange reaction through a generic string search rather than its exact model identifier (*EX_succ_e*), applied knockouts by directly setting reaction bounds rather than using COBRApy’s native *knock_out()* method, and operated without context manager protection against in-place model modification. As a result, the code ran and returned output but was fragile and inconsistent with COBRApy best practices (Accuracy: 3, Completeness: 5, OPS: 3.60) (Figure 2A). With RAG, ChatGEM retrieved the correct reaction identifier, applied knockouts through the appropriate API, and produced verified results, readable code (Accuracy: 4, Completeness: 5, OPS: 4.30) (Figure 2A).

**Figure 2:**
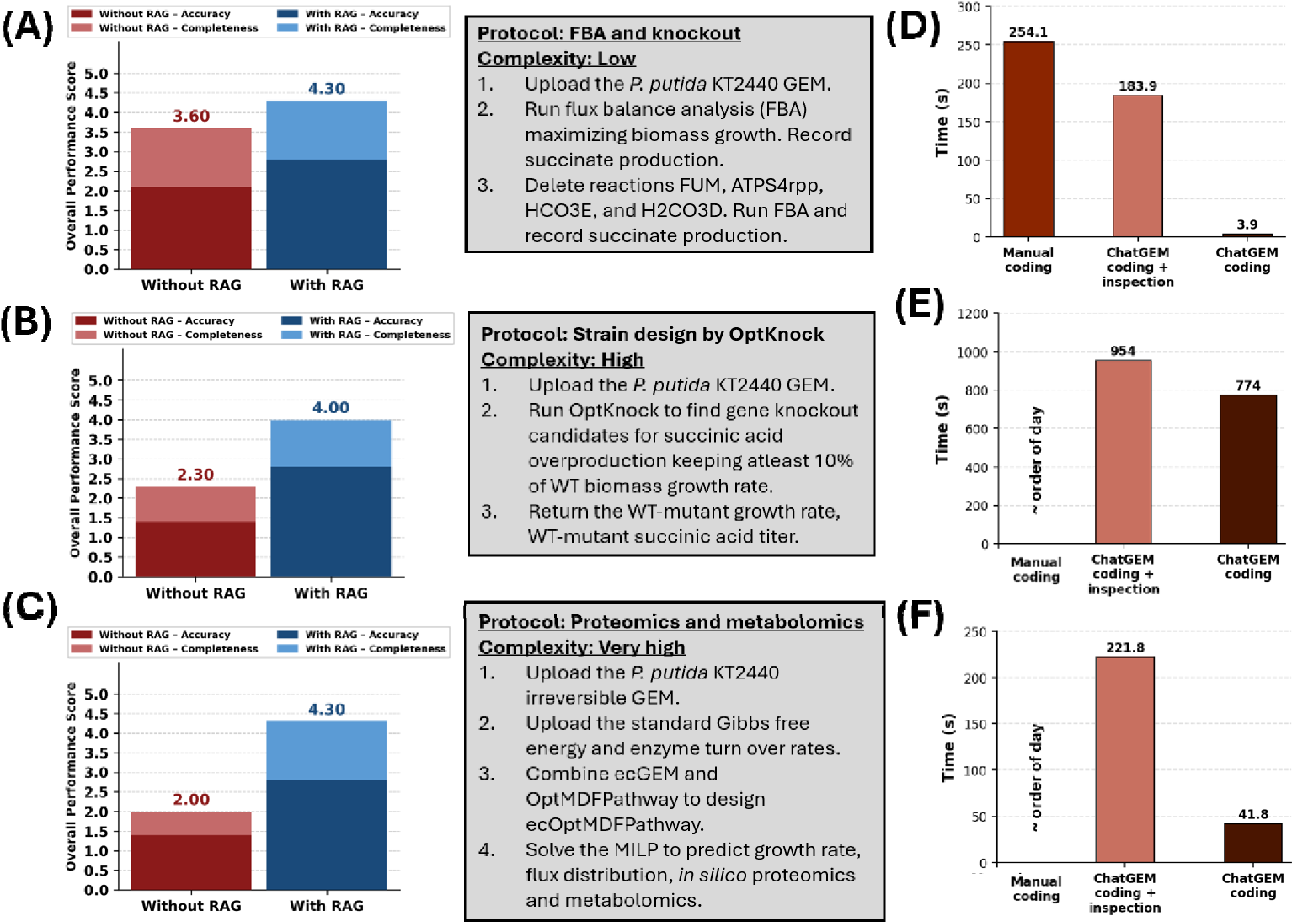
ChatGEM code generation performance and time efficiency across tasks of increasing complexity. (A) Overall Performance Score (OPS) for the FBA and knockout task (low complexity) without RAG and with RAG. The corresponding protocol is shown in the center panel. (B) OPS for the OptKnock strain design task (high complexity) without RAG and with RAG. (C) OPS for the ecOptMDFPathway proteomics and metabolomics integration task (higher complexity) without RAG and with RAG. (D) Time comparison for the FBA and knockout task across manual coding, ChatGEM coding with manual inspection, and ChatGEM coding alone. (E) Time comparison for the OptKnock strain design task across manual coding, ChatGEM coding with manual inspection, and ChatGEM coding alone. (F) Time comparison for the ecOptMDFPathway task across manual coding, ChatGEM coding with manual inspection, and ChatGEM coding alone.

The performance gap widened substantially for the OptKnock strain design task. When running without RAG, ChatGEM used Cameo library (Cardoso, et al., 2018), a reasonable choice in isolation, but a direct violation of the protocol requiring the StrainDesign toolbox. Despite some syntax error and the code being logically structured and complete in its steps, the library mismatch alone rendered it non-compliant (Accuracy: 2, Completeness: 3, OPS: 2.30) (Figure 2B). With RAG, ChatGEM correctly formulated the OptKnock bilevel problem using *sd.OPTKNOCK*, enforced biologically realistic constraints including a glucose uptake rate of -6.0 mmol/gDW/h and a minimum growth requirement of 10% wildtype, excluded nine essential reactions from the knockout search space, and exported a structured summary to Excel (Accuracy: 4, Completeness: 4, OPS: 4.00) (Figure 2B).

The most striking contrast emerged in the ecOptMDFPathway task requiring integrating enzyme-constrained modeling with thermodynamic feasibility analysis. Without RAG, the generated code contained a critical logical error in the stoichiometric mass balance, alongside an incorrect formulation of the thermodynamic driving force constraint, and the code could not execute (Accuracy: 2, Completeness: 2, OPS: 2.00) (Figure 2C). With RAG, ChatGEM assembled a complete and functional MILP enforcing mass balance, thermodynamic feasibility derived from ΔG° values and metabolite concentrations, flux-activity coupling via binary variables, a proteome capacity ceiling of 0.60 g/gDW, and a minimum biomass flux constraint, exporting flux, concentration, thermodynamic, and protein usage results across four structured Excel sheets (Accuracy: 4, Completeness: 5, OPS: 4.30) (Figure 2C).

Beyond code quality, ChatGEM substantially reduced the time required to complete each modeling task. For the FBA and knockout task, manual coding required 254.1 seconds, which ChatGEM reduced to 3.9 seconds when executed without inspection, a 65-fold reduction (Figure 2D). For the OptKnock strain design task, ChatGEM coding alone completed in 774 seconds compared to more than a day for manual coding (Figure 2E). For the ecOptMDFPathway task, ChatGEM coding alone required 41.8 seconds compared to more than a day of manual coding like the OptKnock case (Figure 2F). Notably, even ChatGEM coding with manual inspection consistently requires far less time compared to manual coding alone, reflecting even ChatGEM with human in the loop can also consistently beat manual coding tasks (Figure 2 D-E).

To evaluate whether model choice affects code generation performance, we compared the default o4-mini, a lightweight reasoning model, against GPT-4.1, a general-purpose frontier model with substantially higher inference cost, across the same three tasks under both without-RAG and with-RAG conditions (Supplementary Figure 1; Supplementary File 2). Without RAG, o4-mini achieved a mean OPS of 2.63 compared to 2.18 for GPT-4.1, demonstrating that the reasoning-optimized architecture of o4-mini confers a consistent advantage in the absence of domain-specific context. On the ecOptMDFPathway task, GPT-4.1 produced nonlinear proteome constraint formulations that Gurobi would reject, while o4-mini generated a structurally sounder MILP. With RAG, both models converged to identical mean OPS of 4.20 and matched scores across all three tasks, indicating that RAG grounding effectively equalizes performance between models by supplying the domain-specific knowledge that reasoning capacity alone cannot substitute for. Given that o4-mini matches GPT-4.1 with RAG while outperforming it without RAG, and at a fraction of the inference cost, o4-mini represents the more practical and cost-effective backbone for ChatGEM’s code generation pipeline without sacrificing output quality.

### Case Study: Succinate Overproduction form *Pseudomonas putida* KT2440

To demonstrate ChatGEM’s utility in a realistic metabolic engineering context, we applied it to rank *P. putida* KT2440 strains for succinate overproduction using ecGEM. Four engineered strains were evaluated: wild-type (KT2440), a SAGE-compatible strain containing 3 landing pads for serine recombinase-aided DNA insertion (Landing Pad), a constitutively expressed dCas12a construct (Constitutive), and a crystal-violet inducible dCas12a construct (Induced). Each strain was grown on glucose as the sole carbon substrate with proteomic sampling at 12 and 24 hours alongside direct measurement of succinate secretion.

Proteomics data were processed using intensity-Based Absolute Quantification (iBAQ) protein abundances, and enzyme turnover rates were calculated using TurNuP (Kroll, et al., 2023) to parameterize ecGEMs. These information was the input to ChatGEM, which reconstruction eight ecGEMs in total, spanning four strains across two time points. Model quality was assessed by comparing ecGEM-predicted enzyme requirements against iBAQ-derived enzyme abundances across the proteome. The two showed strong agreement (r = 0.50, Figure 3A), confirming that the models captured realistic enzyme allocation patterns. Predicted growth rates were similarly consistent with biologically plausible ranges across all eight experimental conditions (Figure 3B) as defined in MEMOTE (Kroll, et al., 2023), providing further confidence in model predictions.

**Figure 3:**
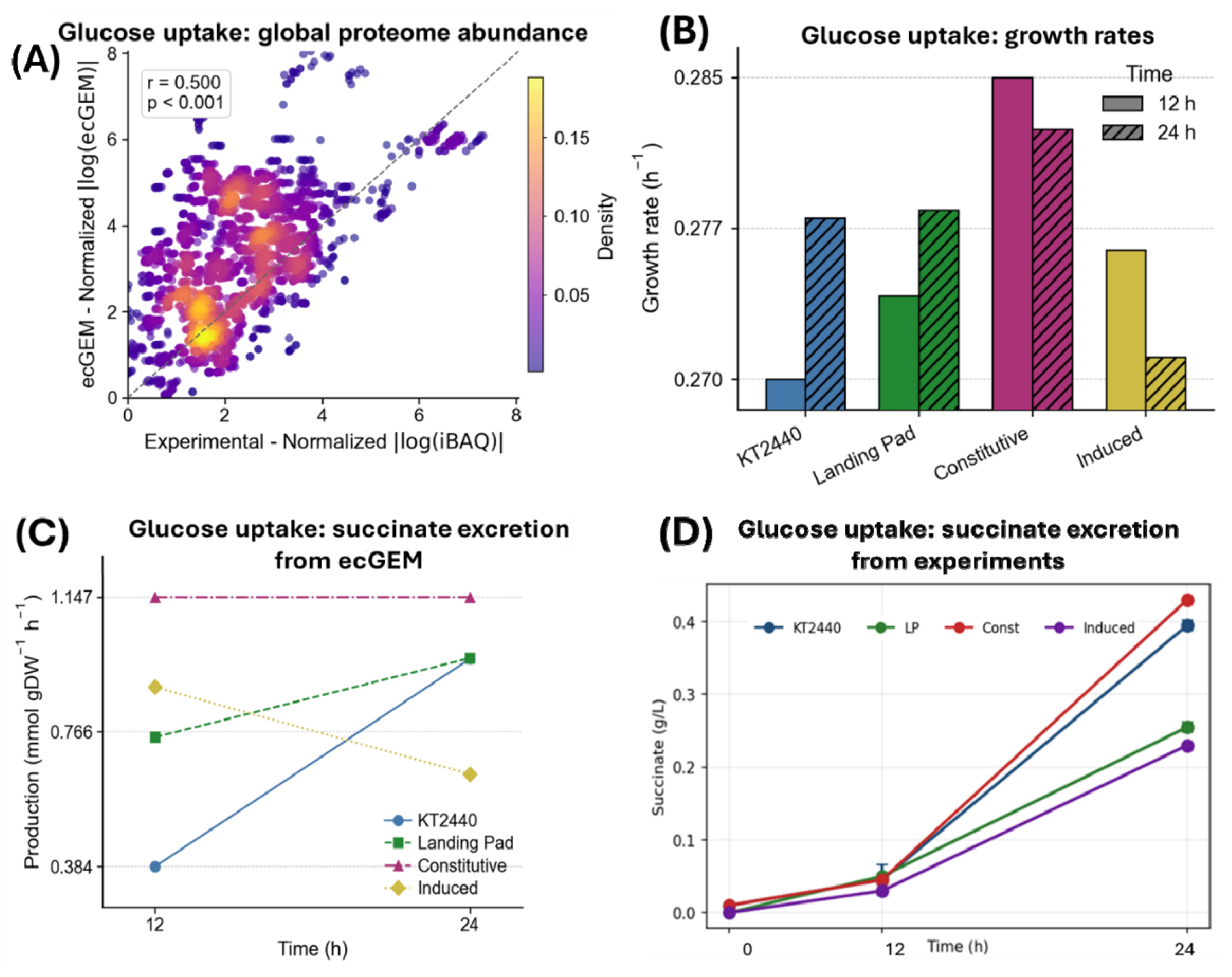
Enzyme-constrained GEM validation and succinate production across *P. putida* KT2440 strains grown on glucose. (A) Correlation between experimentally measured normalized iBAQ protein abundances and ecGEM-predicted normalized enzyme requirements across the global proteome under glucose uptake conditions. Density of data points is indicated by the color scale. (B) ecGEM-predicted growth rates for KT2440, SAGE Landing Pad, constitutive dCas12a, and inducible dCas12a strains at 12 h (solid bars) and 24 h (hatched bars) under glucose uptake conditions. (C) ecGEM-predicted succinate excretion rates for all four strains at 12 and 24 hours, showing a general increase over time except for the Induced strain. (D) Experimentally measured succinate titers over 24 hours for KT2440, Landing Pad (LP), Constitutive (Const), and Induced strains, consistent with ecGEM predictions in panel C.

Analysis of ecGEM-predicted succinate fluxes revealed divergent production trajectories across the four strains from 12 to 24 hours (Figure 3C). KT2440 and Landing Pad showed a consistent increase over time, a trend mirrored closely by experimentally measured succinate titers (Figure 3D). The Constitutive strain maintained the highest succinate excretion rate across both time points, remaining essentially flat from 12 to 24 hours. The Induced strain was the sole exception to any upward or flat trend, declining from 12 to 24 hours in simulation in ecGEM, suggesting a strain-specific regulatory response that the model was unable to capture therefore requiring further investigation. To identify the metabolic basis for differential succinate production across strains, we examined enzyme usage across the central carbon pathway for different reactions (Figure 4 A-B). EDD emerged as the largest consumer of the total enzyme pool, followed by GLCDpp and ACONTa, highlighting the Entner-Doudoroff pathway as a dominant flux route and a potential bottleneck for carbon redirection toward succinate, consistent with literature (Nikel, et al., 2015).

**Figure 4.**
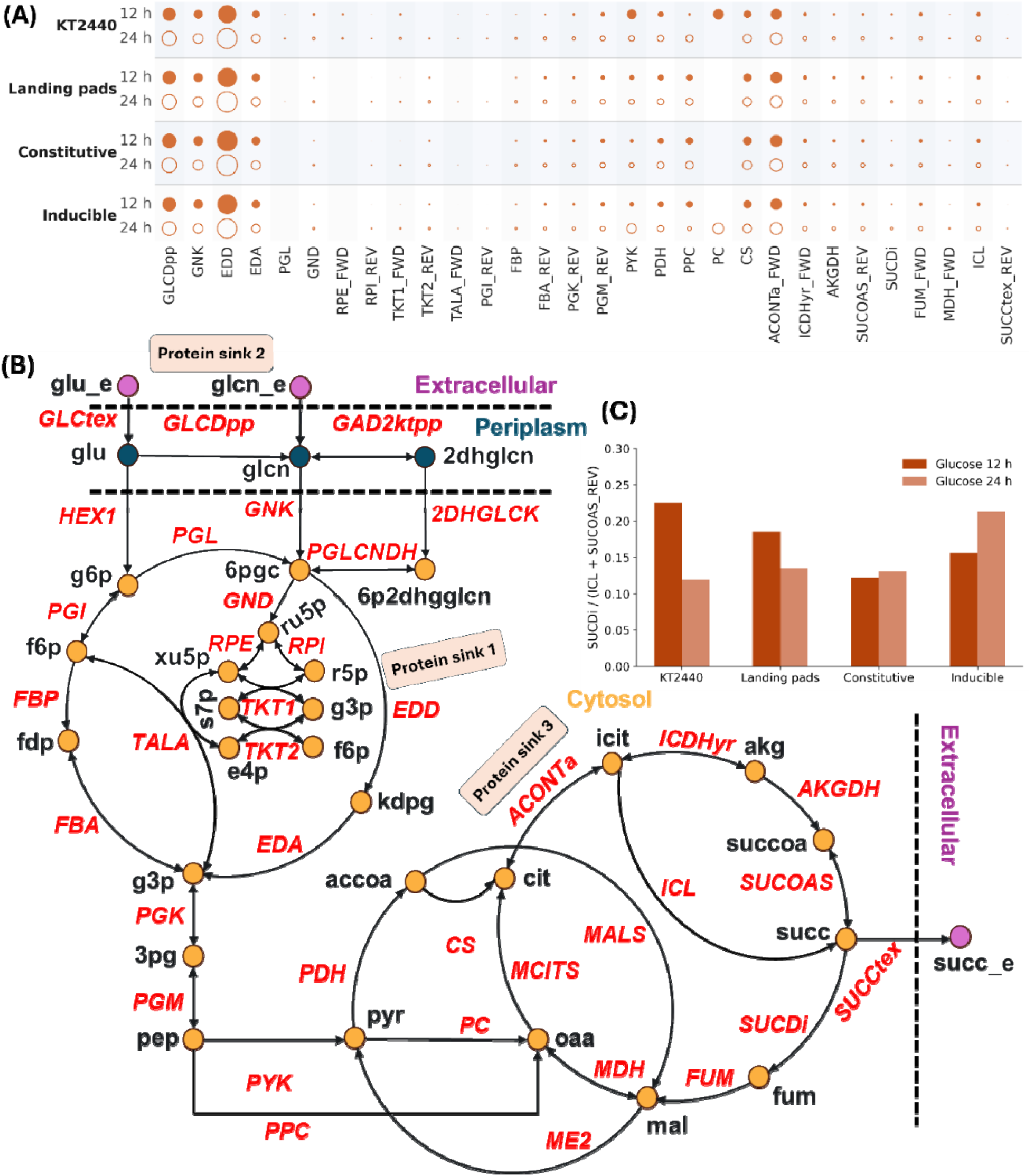
Central carbon pathway enzyme allocation and succinate leakage index across *P. putida* KT2440 strains. (A) Bubble plot of ecGEM-predicted enzyme usage across central carbon pathway reactions for KT2440, Landing Pad, Constitutive, and Inducible strains at 12 h and 24 h under glucose uptake conditions. Bubble size reflects the proportion of total enzyme pool allocated to each reaction. EDD, GLCDpp, and ACONTa emerge as the dominant protein sinks across all strains. (B) Map of the central carbon metabolic network in *P. putida* KT2440, with reactions colored in red and metabolites a nodes. Three major protein sinks are annotated: EDD (Protein sink 1), GLCDpp (Protein sink 2), and ACONTa (Protein sink 3). (C) SLI for all four strains at 12 h (dark orange) and 24 h (light orange) under glucose uptake conditions. The Constitutive strain exhibits the lowest SLI at both time points (0.12 and 0.13), indicating minimal enzyme allocation toward succinate consumption and identifying it as the most favorable chassis for succinate overproduction engineering.

We defined the ratio of enzyme usage allocated to SUCDi (succinate dehydrogenase) relative to the sum of ICL (isocitrate lyase) and SUCOAS (succinyl-CoA synthetase) as succinate leakage index (SLI), a quantitative metric for strain engineering potential (Figure 4B). A high SLI indicates that a greater proportion of the enzyme pool is directed toward consuming succinate rather than producing it, reducing the strain’s amenability to further engineering. Ranking strains by this index identified the Constitutive strain as the most promising chassis, with the lowest SLI of 0.12 and 0.13 at 12 and 24 hours, respectively (Figure 4C). By contrast, the WT strain exhibited the highest SLI at 12 hours (0.22), reflecting substantial enzyme allocation toward succinate consumption early in growth, which decreased to 0.12 by 24 hours. The Landing Pad strain showed intermediate values of 0.19 and 0.13 at 12 and 24 hours. Notably, the Inducible strain was the only strain whose SLI increased over time, rising from 0.16 at 12 hours to 0.21 at 24 hours, consistent with its divergent succinate production trajectory observed in the ecGEM predictions. The SLI ranking agrees with experimental succinate titers, which were highest in the Constitutive strain across both time points (Figure 3D), suggesting that minimizing enzyme allocation toward succinate oxidation is a reliable and experimentally consistent criterion for strain prioritization.

## DISCUSSION

GEMs have long been considered as a powerful tool in biology to quantify metabolism, yet their adoption outside computational groups has remained constrained by the stiff programming barrier. ChatGEM addresses this directly by embedding the full GEM simulation stack within a conversational multi-agent interface, enabling researchers to formulate and execute complex GEM problems through natural language. RAG-grounded code generation produced a 60% improvement in OPS, defined earlier, relative to unaided generation.

The evaluation framework used here, based on accuracy and completeness combined into a weighted OPS, is consistent with approach adopted in a recent effort, BioAgents (Mehandru, et al., 2025). BioAgents, a RAG-enhanced multi-agent system for bioinformatics workflow generation, similarly evaluated performance on these axes across tasks of increasing complexity, finding that systems matched human expert performance on conceptual tasks but degraded as code generation complexity increased, a pattern parallel to ChatGEM’s without-RAG condition. ChatGEM’s RAG architecture directly addresses this failure mode by grounding generation in verified, domain-specific documentation, maintaining a consistent performance floor even at the varying complexity tier. To evaluate whether model choice affects this pattern, we compared o4-mini, a lightweight reasoning model, against GPT-4.1, a general-purpose frontier model with substantially higher inference cost, across all three tasks under both conditions (Figure 2 A-C; Supplementary Figure 1; Supplementary File 1; Supplementary File 2). Without RAG, o4-mini achieved a mean OPS of 2.63 versus 2.18 for GPT-4.1. With RAG, both models converged to an identical mean OPS of 4.20 across all three tasks, confirming that RAG grounding effectively equalizes performance by supplying the domain-specific knowledge that reasoning capacity alone cannot substitute for. Given that o4-mini matches GPT-4.1 with RAG while outperforming it without, o4-mini represents the more cost-effective agent for ChatGEM. This advantage of reasoning models over general-purpose models on structured, multi-step tasks is well documented. For example, on the BIG-Bench Extra Hard benchmark, the best reasoning-specialized model achieved an accuracy of 54.2% compared to 23.9% for the best general-purpose model, with the largest performance gaps concentrated in tasks requiring algorithmic planning and multi-step logic (Kazemi, et al., 2025). The pattern observed in ChatGEM, where o4-mini outperforms GPT-4.1 without RAG but converges with it under RAG grounding, suggests that reasoning capacity and domain-specific context retrieval are partially substitutable, with RAG effectively compensating for the absence of deep reasoning in general-purpose models on well-defined coding tasks.

The emergence of LLM-based tools has begun reshaping how biologists interact with complex data. Natural language interfaces now span single-cell transcriptomics, where CellWhisperer (Schaefer, et al., 2025) leveraged multimodal contrastive learning across over one million transcriptomes to make RNA-seq exploration conversational. In metabolic engineering, D2Cell (Li, et al., 2026) distilled over 29,000 literature entries into a structured knowledge base, pairing it with a hybrid deep learning and GEM-based model for target prediction. For GEM reconstruction, it is demonstrated that LLM-assisted curation produced Human2, a high-quality consensus human GEM, resolved to tissue and organ specificity (Luo, et al., 2026). ChatGEM extends this trajectory into the domain of GEM-based analysis and by only using natural language commands. The common thread is not automation of expertise but its amplification, with LLMs serving as executable interfaces for knowledge that would otherwise remain inaccessible.

The biological case study reinforces this point. Applied to four *P. putida* KT2440 strains, ChatGEM coordinated the reconstruction of eight accurate ecGEMs, a task that conventionally requires deep familiarity with the enzyme-constrained modeling and manual reconciliation of proteomics abundance data with enzyme kinetic parameters (Chen, et al., 2024; Domenzain, et al., 2022). The ecGEM-predicted proteome correlated with iBAQ-derived abundances at r = 0.50 (p < 0.001), comparable to agreement reported for *P. putida* models constrained with proteomic and kinetic data (Bujdos, et al., 2023). ChatGEM identified EDD, GLCDpp, and ACONTa as dominant protein sinks, consistent with the known dominance of the Entner-Doudoroff pathway in *P. putida* (Nikel, et al., 2015). The Constitutive strain, with the lowest SLI at both time points (0.12 and 0.13), was predicted as the optimal chassis and observed by experimental succinate titers. ChatGEM delivered this complete analysis through natural language represents a capability not previously demonstrated in any conversational AI framework for metabolic engineering, to the best of our knowledge.

Finally, ChatGEM represents a qualitative shift in how GEM can be practiced. While existing LLM-based tools have advanced natural language interfaces for transcriptomics exploration and literature-driven target prediction, ChatGEM operates at the level of quantitative mechanistic simulation, executing constraint-based modeling workflows, strain design, ecGEM reconstruction, and derivation of experimentally observed engineering metrics, all through conversation. The ability to execute these workflows through natural language interaction addresses a long-standing bottleneck in metabolic engineering, where computational expertise has historically constrained who can engage with GEM-based design.

## MATERIALS AND METHODS

### ChatGEM implementation and workflow

ChatGEM was developed using ADEPT as the reference architecture. We used infrastructure-as-code to deploy and test ADEPT on AWS EC2 instances as well as on an internal on-site high-performance computing system. The baseline ADEPT framework was adapted to support a custom Python code execution environment built with additional dependencies for COBRApy, StrainDesign, and Gurobi. In addition, we built a custom Gurobi license manager tool to support internal licensing requirements. Finally, we adjusted the execution code and run-time allowances to ensure long-running jobs (i.e. multi-knockout strain design) would be complete. Detailed documentation on how to deploy and customize ADEPT are available at https://github.com/pnnl/adept-agentic.

Once deployed, we primarily utilized ADEPT’s built-in streamlit app which connects to the persistent ADEPT services and provides a chat interface and on-demand file ingestion. A set of 52 human curated python scripts were then uploaded and processed for RAG via the interface. These scripts include examples ranging from simple flux balance analysis to ecOptMDFPathway (see Data Availability). During a typical workflow, a user would then ask a query in the chat interface, triggering ADEPT’s Scientific Workflow Agent. Any code execution by the agent was routed to the custom ChatGEM sandbox container, which provided a secure environment to perform *in silico* metabolic engineering. Finally, the outputs were sent to a separate, standalone LLM-as-judge (Claude Sonnet 4.6) for OPS scoring.

### *P. putida* strains construction Strain construction

Wild-type *P. putida* KT2440 or a variant of *Pseudomonas putida* KT2440 that is compatible with Serine recombinase-Assisted Genome Engineering (SAGE) called AG5577 were used for this study (Elmore, et al., 2023). AG5577 has three separate poly-*attB* sequences incorporated into its genome (Huenemann, et al., 2025). The dCas12a expression plasmids were incorporated into AG5577 using SAGE as described in detail previously (*ref SAGE). In brief, 50 µL of electrocompetent *P. putida* AG5577 cells were mixed with 200 ng of the non-replicating TG1 recombinase expression plasmid pGW38 (Elmore, et al., 2023) and 200 ng of non-replicating target plasmid at room temperature. The target plasmids were either pJE1778 (constitutive dCas12a) or pJE2165 (crystal violet inducible dCas12a). Cells were electroporated, resuspended in 950 µL SOC (BD), recovered at 30 C for 1 hr, and plated onto solid selective LB medium (VWR) containing 30 µg/mL gentamicin sulfate (Sigma Aldrich) and 15 g/L agar. Cells were incubated overnight at 30 °C. Gentamicin resistant colonies were generated as a consequence of recombination between the TG1 *attB* sequence in the AG5577 chromosome and a TG1 *attP* sequence in the target plasmid. One colony from each transformation was selected for further use.

### Plasmid construction

Q5® High-Fidelity DNA Polymerase (New England Biolabs - NEB) and primers synthesized by Eurofins Genomics were used in all PCR amplifications for plasmid construction. OneTaq® DNA polymerase (NEB) was used for colony PCR. EvaGreen® dye (Biotium) was supplemented at 1x final concentration in PCR reactions for qPCR applications. Plasmids were constructed by Gibson Assembly using NEBuilder® HiFi DNA Assembly Master Mix (NEB). Plasmids were transformed into chemically competent NEB 5-alpha F’I^q^ (NEB). Standard chemically competent *E. coli* transformation protocols were used to construct plasmid host strains. Transformants were selected on LB (Lennox) agar plates containing antibiotics (30 µg/mL gentamicin sulfate) for selection and incubated at 37 °C. Template DNA for PCR was synthesized by Twist Biosciences. Zymoclean Gel DNA recovery kit (Zymo Research) was used for all DNA gel purifications. Plasmid DNA was purified from *E. coli* using GeneJet plasmid miniprep kit (ThermoScientific) or ZymoPURE II plasmid midiprep kit (Zymo Research). The sequence of plasmid pJE2165 was confirmed using Oxford Nanopore sequencing by Plasmidsaurus. Plasmids and strains used in this work are listed in Table 1, and maps of plasmids constructed in this study can be found in Supplementary File 4.

**Table 1.**
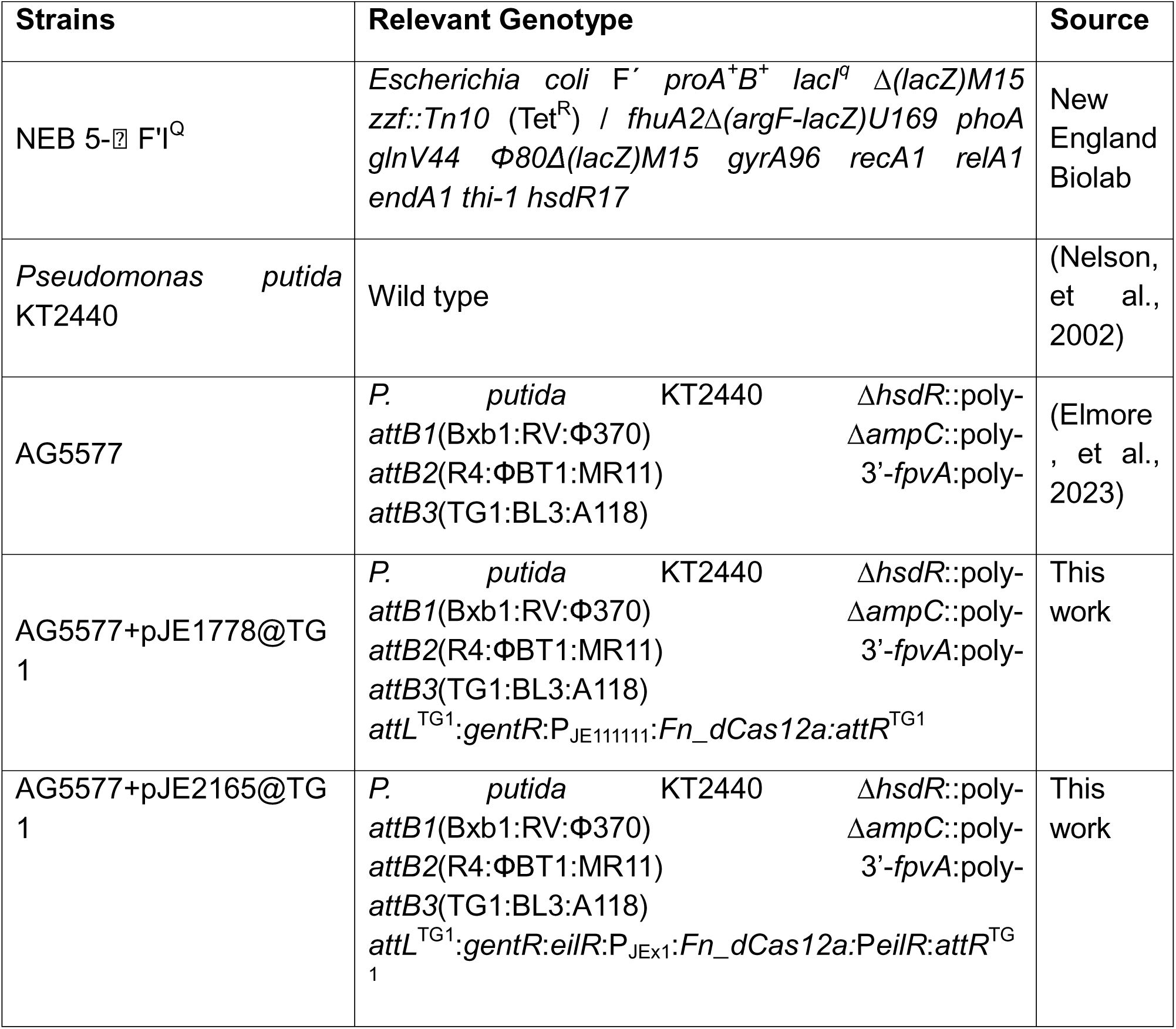
Strains and plasmids used in this work.

### Growth conditions of strains and sample preparation for proteomics data

Media. P. putida strains were grown on LB media or agar and a variation of M9 minimal media previously described. In addition to the standard formula, 30 mM MOPS was added to provide buffering at pH 7.4. A nitrogen limited version of the M9 media was utilized for production with 0.28 g/L of (NH4)2SO4 instead of the standard concentration.

For cultivation, *P. putida* strains were initially streaked from cold storage on LB agar plates containing appropriate antibiotic concentrations for the respective strains (gentamycin: 15 µg/ml). Single colonies were used to inoculate 35 mL of LB media in a 125 mL unbaffled shaking flask and grown to saturation (∼24 h). LB seed cultures were then used to inoculate the standard nitrogen M9 minimal media formula and grown overnight (∼12-14 h). The M9 seed was then used to inoculate the nitrogen limited M9 media at an initial OD600 of 0.1. Strains were cultivated in 24 well plates containing 2 mL of media, sealed with a breathable seal and without lids. (Incubator shaking). At 6, 12, or 24 h a single plate containing four replicates were sampled and sacrificed for HPLC and proteomic samplings. Cultivations occurred at 30 °C.

HPLC samples were filtered through 0.2 µM filter and ran according to the following procedure. Filtered samples were analyzed by HPLC on a system fitted with a Waters 2489 UV/Visible detector reading absorbance at 210 nm. Separation was performed on a Bio-Rad Aminex HPX-87 H ion exclusion column (300 mm x 7.8 mm) held at 50 °C, with 0.0045 M sulfuric acid as the mobile phase at a flow rate of 0.55 mL/min. Metabolite concentrations were determined from linear regression calibration curves generated for each analyte (Czajka, et al., 2025). Proteomic samples were centrifuged at 5,500 x g for 2 mins, washed with 4 °C sterile PBS, pelleted, and flash frozen in liquid nitrogen. Samples were stored at –80 °C until processing.

### Proteomics data analysis

Proteomics data were obtained from four *P. putida* KT2440 strains (WT, Landing Pad, Constitutive, and Inducible) grown on glucose as the sole carbon source, with samples collected at 12 and 24 hours in biological triplicate. Proteins were digested following the previously described sample preparation protocol (Gao, et al., 2020) prior to LC-MS/MS analysis. LC MS/MS data were acquired on a Thermo Vanquish Neo LC system configured in a trap and elute setup and coupled to an Thermo Orbitrap Astral mass spectrometer. Samples at 0.1 µg/µL were injected at 5 µL and separated using mobile phase A (water + 0.1% formic acid) and mobile phase B (acetonitrile + 0.1% formic acid). Peptides were first loaded onto a PepMap Neo Trap Cartridge at 200 µL/min, then resolved on an ES906 PepMap analytical column (150 mm × 150 µm ID, C18, 2 µm, 100 Å) maintained at 45 °C. The LC gradient used variable flow, with separation performed at 0.8 µL/min from 3.0 to 23.0 min while increasing from 8% to 45% B, followed by a high organic wash at 99% B and re equilibration to 1% B. The total LC method time was 25.7 min.

Mass spectrometry data were acquired using a data independent acquisition (DIA) method with interleaved full MS1 and DIA MS2 scans. Full MS1 scans were acquired with 240,000 resolution power, precursor range of 380–980 m/z, and AGC target of 5e6. DIA-MS2 scans were collected using 25% HCD collision energy and AGC target of 5e4, with contiguous 2 m/z isolation windows spanning 380–980 m/z. DIA data were processed using DIA NN 2.3.1 Academia (Demichev, et al., 2020). Searches used a DIA NN–predicted library for Pseudomonas putida KT2440 supplemented with heterologous dCas12a and gentamicin resistance protein sequences. Match between runs/reanalysis, smart profiling, relaxed protein inference, heuristic protein grouping, and gene level protein grouping were enabled. In-silico digestion was performed at K/R sites with N terminal methionine excision allowed. Carbamidomethylation of cysteine was set as a fixed modification, while methionine oxidation and N terminal acetylation were set as variable modifications, with up to three variable modifications per precursor permitted. Search results were filtered using the following criteria: Q.Value ≤ 0.01, Lib.Q.Value ≤ 0.01, PG.Q.Value ≤ 0.01, Lib.PG.Q.Value ≤ 0.01, Quantity.Quality ≥ 0.5, and PG.MaxLFQ.Quality ≥ 0.7.Raw precursor intensities were imported from DIA-NN output and processed using the pmartR (Stratton, et al., 2024) package in R. Peptide-level data were structured with precursor identifiers as features, sample identifiers as observations, and protein group annotations from the *P. putida* KT2440 UniProt reference proteome (2017-04-07) as metadata. Missing values were represented as NA and data were transformed to the log2 scale. Samples were grouped according to combinations of strain, carbon source, and time point. In preparation for statistical comparisons using IMD-ANOVA (Webb-Robertson, et al., 2010), any peptides not meeting a minimum number of observations per group were excluded from the dataset (minimum of two non-missing observations per group for ANOVA and three for the g-test). Potential sample outliers were identified using the robust Mahalanobis distance (RMD) filter (Matzke, et al., 2011) based on four metrics: median absolute deviation, skewness, pairwise correlation, and proportion of missing values. All samples were normalized by global median centering after verifying the lack of dependence between sample medians and group using the Kruskal-Wallis test (p = 0.49) (Webb-Robertson, et al., 2010).

Statistical analyses at both the peptide and protein level were performed as follows. Peptides were rolled up to the protein level using a reference-based rollup method available in the pmartR package (“rrollup”), in which peptides are first scaled by a reference peptide (the one with the least missingness) and protein abundance is set as the median of these scaled peptides. For quantitative comparisons of log2-fold change at both the peptide- and protein-level, Holm-adjusted two-sample t-tests were used.

Quantitative comparisons were made using Holm-adjusted G-tests, which assess systematic missingness between groups. The Holm adjustments were applied within each peptide (protein) and separately to the following sets of comparison groups: strain comparisons - 24 comparisons per peptide (protein), time point comparisons - 8 comparisons per peptide (protein), carbon source comparisons - 8 comparisons per peptide (protein)

### iBAQ implementation on proteomics data

Protein-level iBAQ intensities were computed following the previously described approach (Sanchez, et al., 2017). After the filtering and normalization described above, peptide-level intensities were first consolidated to the protein level using MaxLFQ via the iq (Pham, et al., 2020) package. The resulting protein-level intensities were back-transformed to linear scale and divided by the number of theoretically observable tryptic peptides per protein, defined as peptides of length 7 to 30 amino acids generated by in silico tryptic digestion of the reference FASTA (cleavage after K or R, not before P) using the *P. putida* KT2440 proteome sequence file (ID_008893_17C184F0). For protein groups with multiple members, the leading protein was defined as the member with the highest molecular weight, and the corresponding tryptic peptide count was used for iBAQ normalization. Final iBAQ values were organized into a wide-format matrix indexed by protein group and exported for downstream ecGEM integration.

### Enzyme constrained metabolic modeling

Enzyme-constrained genome-scale metabolic models were reconstructed for four *P. putida* KT2440 strains at two time points (12 h and 24 h) under glucose uptake conditions, yielding eight models in total. The irreversible formulation of iJN1463 was used as the base model with BIOMASS_KT2440_WT3 as the objective function, and all optimization was performed using COBRApy and Gurobi solver.

For each reaction in the model, the molecular weight of the catalyzing enzyme was computed from its amino acid sequence retrieved from the *P. putida* KT2440 UniProt reference proteome. Enzyme turnover numbers were retrieved from TurNuP, a machine learning model that predicts values from sequence and structural features (Kroll, et al., 2023). Reaction-specific upper bounds were then computed by combining iBAQ-derived enzyme abundances with TurNuP values, following the capacity constraint formulation (Sanchez, et al., 2017). Upper bounds were applied to 2,479 reactions with matched proteomics coverage; reactions without coverage retained their stoichiometric model upper bounds. A global proteome capacity constraint of 0.20 g gDW⁻¹ was imposed across all models, consistent with the total measured protein content of *P. putida* KT2440 under glucose growth conditions.

For each condition, an unconstrained model was first solved to identify reactions whose iBAQ-derived upper bounds were violated relative to the unconstrained optimal flux by more than 100%, indicating cases where proteomics measurement uncertainty or incomplete coverage would otherwise render the model infeasible. Upper bounds for these reactions were relaxed to 1,000 mmol gDW⁻¹ h⁻¹ prior to final model solution. Each solved ecGEM produced a full flux distribution and per-reaction protein usage estimate, used for downstream analysis of central carbon pathway enzyme allocation and computation of the succinate leakage index.

## SUPPLEMENTARY FILES

Supplementary Figure 1: Comparison between o4-mini and ChatGPT 4.1.

Supplementary File 1: Evaluation of codes generated from ChatGEM (with and without RAG).

Supplementary File 2: o4-mini vs ChatGPT 4.1 generated code and reports.

Supplementary File 3: Proteomics Data for the used conditions.

Supplementary File 4: Maps of plasmids constructed in this study

## DATA AVAILABILITY

The codebase for RAG, eight different ecGEMs, and iBAQ implementation codes are available at https://github.com/pnnl/ChatGEM. ADEPT is available at https://github.com/pnnl/adept-agentic.

## AI USAGE STATEMENT

Artificial intelligence tools were used to assist with editorial refinement of selected portions of the manuscript text. All scientific content, experimental design, data collection, computational analyses, and interpretations are the original work of the authors. AI assistance was limited to language editing, generation of Figure 1, and did not contribute to the conception, execution, or conclusions of the research presented.

## ACKNOWLEDGEMENT

This research was conducted as part of the Predictive Phenomics Initiative at Pacific Northwest National Laboratory (PNNL) and was supported by the Laboratory Directed Research and Development (LDRD) Program at PNNL. The development of ADEPT was also supported by the Generative AI Investment, conducted under the LDRD Program at PNNL. This work was further supported by the Orchestrated Platform for Autonomous Laboratories (OPAL) project, funded by the U.S. Department of Energy (DOE), Office of Science, Office of Biological and Environmental Research, and by the Office of Science, Office of Advanced Scientific Computing Research, under FWP 86481. We thank Adam Guss at Oak Ridge National Laboratory for kindly sharing *P. putida* AG5577. Pacific Northwest National Laboratory is a multi-program national laboratory operated by Battelle for the U.S. Department of Energy under Contract No. DE-AC05-76RL01830.

