## Supplementary material for "ChatGEM: An Agentic Architecture Enabling Interactive Simulation of Genome-Scale Metabolic Models": Generated code was evaluated by Claude Sonnet 4.6 with regards to two metrics (Supplementary File 1) (from another version of this preprint: version 1, 2026-07-21)

**Model: o4-mini**

**Task1: FBA and knockout; Complexity: Low**

**Protocol:**

I have uploaded the iJN1463.xml model of *Pseudomonas putida* KT2440. Please perform the following using the codebase and Gurobi solver:

A) Run baseline FBA and record biomass growth rate and succinate exchange flux.

B) Knock out reactions: FUM, ATPS4rpp, HCO3E, H2CO3D.

C) Rerun FBA after knockouts.

D) Report biomass and succinate exchange flux before and after deletions

**Without RAG Code:**

#!/usr/bin/env python3

import cobra

import os

def find_model(path, name_substr):

"""

Recursively search for a file containing name_substr under path.

Returns full filepath or raises FileNotFoundError.

"""

for root, dirs, files in os.walk(path):

for f in files:

if name_substr.lower() in f.lower() and f.lower().endswith('.xml'):

return os.path.join(root, f)

raise FileNotFoundError(f"No SBML XML file containing '{name_substr}' found under {path}")

def run_fba(model, succ_regex="succ"):

"""

Optimize model and return objective value and succinate exchange flux.

We identify the first EX_ reaction whose ID or name contains succ_regex.

"""

sol = model.optimize()

biomass = float(sol.objective_value)

\# find succinate exchange reaction

succ_rxn = None

for rxn in model.exchanges:

if succ_regex in rxn.id.lower() or succ_regex in (rxn.name or "").lower():

succ_rxn = rxn

break

succ_flux = float(sol.fluxes[succ_rxn.id]) if succ_rxn else None

return biomass, succ_flux

def main():

\# 1) Locate the SBML file

upload_dir = "/app/data/uploaded_files"

sbml_path = find_model(upload_dir, "ijn1463")

\# 2) Load model and set Gurobi

model = cobra.io.read_sbml_model(sbml_path)

try:

model.solver = "gurobi"

except Exception:

print("Warning: Could not set solver to gurobi, using default solver:", model.solver)

\# 3) Baseline FBA

baseline_biomass, baseline_succ = run_fba(model)

print(f"Baseline biomass	= {baseline_biomass:.6f} h^-1")

print(f"Baseline succinate ex = {baseline_succ:.6f} mmol·gDW^-1·h^-1")

\# 4) Knock out reactions

to_knock = ['FUM', 'ATPS4rpp', 'HCO3E', 'H2CO3D']

for rid in to_knock:

if rid in model.reactions:

model.reactions.get_by_id(rid).bounds = (0.0, 0.0)

print(f"Knocked out reaction {rid}")

else:

print(f"Warning: reaction {rid} not found in model")

\# 5) Post‐knockout FBA

post_biomass, post_succ = run_fba(model)

print(f"\nPost-knockout biomass	= {post_biomass:.6f} h^-1")

print(f"Post-knockout succinate ex = {post_succ:.6f} mmol·gDW^-1·h^-1")

if __name__ == "__main__":

main()

**With RAG Code:**

from cobra.io import read_sbml_model

model = read_sbml_model("iJN1463.xml")

model.solver = "gurobi"

sol = model.optimize()

print("Before")

print("Biomass =", sol.objective_value)

print("Succinate =", sol.fluxes["EX_succ_e"])

for rxn in ["FUM", "ATPS4rpp", "HCO3E", "H2CO3D"]:

model.reactions.get_by_id(rxn).knock_out()

sol = model.optimize()

print("\nAfter")

print("Biomass =", sol.objective_value)

print("Succinate =", sol.fluxes["EX_succ_e"])

**Task 2: Strain design by OptKnock, Complexity: High**

**Protocol:**

I have uploaded the iJN1463.xml model of *Pseudomonas putida* KT2440. Please perform the following task using the codebase with StrainDesign toolbox and Gurobi solver:

A) Set glucose uptake to -6.0 mmol/gDW/h.

B) Set the following exchanges to secretion-only (lower bound = 0): EX_co2_e, EX_glcn_e, EX_25dkglcn_e, EX_catechol_e, EX_lac__D_e, EX_meoh_e, EX_4oxptn_e, EX_ac_e.

C) Set biomass objective to BIOMASS_KT2440_WT3 and target to EX_succ_e.

D) Run baseline FBA and record wild-type biomass growth rate and succinate exchange flux.

E) Calculate theoretical maximum succinate by optimizing EX_succ_e directly.

F) Run OptKnock with maximum 2 knockouts and 1 solution, maximizing succinate (outer objective) while maintaining at least 10% wild-type growth (inner objective).

G) Exclude reactions ATPM, CAT, CYO1_KT, CYTBO3_4pp, CYTCAA3pp, NADH16pp, NAt3_1p5pp, PItex, PPK from knockout candidates.

H) Select the first valid solution not containing excluded reactions.

I) Apply knockouts to a mutant model and simulate growth.

J) Report wild-type vs mutant biomass and succinate flux, knockout reactions, and theoretical maximum succinate.

K) Export summary, knockout list, and full flux comparison to OptKnock_iJN1463_succinate.xlsx.

**Without RAG Code:**

#!/usr/bin/env python3

import os

import cobra

import pandas as pd

\# (Optional) if you have Cameo installed for OptKnock

try:

from cameo.strain_design.heuristic import OptKnock

except ImportError:

OptKnock = None

def find_sbml(path, name_substr):

for root, dirs, files in os.walk(path):

for f in files:

if name_substr.lower() in f.lower() and f.endswith('.xml'):

return os.path.join(root, f)

raise FileNotFoundError(f"No SBML .xml file containing '{name_substr}' under {path}")

def run_fba(model, objective=None):

if objective is not None:

model.objective = objective

sol = model.optimize()

return float(sol.objective_value), sol

def main():

#─── A) Load model and set solver ────────────────────────────────────────

upload_dir = "/app/data/uploaded_files"

sbml = find_sbml(upload_dir, "ijn1463")

model = cobra.io.read_sbml_model(sbml)

\# use Gurobi if available

try:

model.solver = "gurobi"

except Exception:

print("WARNING: Gurobi not available, using default solver:", model.solver)

#─── A) Set glucose uptake to -6.0 ──────────────────────────────────────

glc_rxn = model.reactions.get_by_id("EX_glc__D_e")

glc_rxn.lower_bound = -6.0

#─── B) Force secretion‐only for specified exchanges ─────────────────────

to_secrete = [

"EX_co2_e", "EX_glcn_e", "EX_25dkglcn_e", "EX_catechol_e",

"EX_lac__D_e", "EX_meoh_e", "EX_4oxptn_e", "EX_ac_e"

]

for ex in to_secrete:

if ex in model.reactions:

rxn = model.reactions.get_by_id(ex)

rxn.upper_bound = max(rxn.upper_bound, 1000.0) # allow secretion

rxn.lower_bound = 0.0

else:

print(f"WARNING: reaction {ex} not in model")

#─── C) Set biomass objective and target reaction ───────────────────────

biomass_rxn_id = "BIOMASS_KT2440_WT3"

target_rxn_id = "EX_succ_e"

model.objective = biomass_rxn_id

#─── D) Baseline FBA ────────────────────────────────────────────────────

wild_growth, wild_sol = run_fba(model)

wild_succ = float(wild_sol.fluxes[target_rxn_id])

print(f"WILD‐TYPE: growth = {wild_growth:.6f}, succ_ex = {wild_succ:.6f}")

#─── E) Theoretical max succinate ──────────────────────────────────────

max_succ_growth, succ_sol = run_fba(model, objective=target_rxn_id)

max_succ = max_succ_growth

print(f"THEORETICAL MAX succinate = {max_succ:.6f}")

\# Restore biomass objective

model.objective = biomass_rxn_id

#─── F–H) OptKnock design ───────────────────────────────────────────────

\# minimum acceptable growth = 10% of wild‐type

min_growth = wild_growth * 0.10

excluded = [

"ATPM", "CAT", "CYO1_KT", "CYTBO3_4pp", "CYTCAA3pp",

"NADH16pp", "NAt3_1p5pp", "PItex", "PPK"

]

if OptKnock is None:

raise RuntimeError("Cameo not installed: cannot run OptKnock. "

"Install cameo or implement your own MILP.")

ok = OptKnock(

model=model,

biomass=biomass_rxn_id,

target=target_rxn_id,

max_knockouts=2,

minimum_biomass=min_growth,

number_of_solutions=1,

excluded_reactions=excluded

)

result = ok.run() # returns a Cameo StrainDesignResult

ko_list = result.data_frame["knockout_list"][0]

print("Selected knockouts:", ko_list)

#─── I) Apply knockouts and simulate mutant ─────────────────────────────

mutant = model.copy() # copy original model with bounds & uptake set

for rid in ko_list:

mutant.reactions.get_by_id(rid).bounds = (0.0, 0.0)

mut_growth, mut_sol = run_fba(mutant)

mut_succ = float(mut_sol.fluxes[target_rxn_id])

print(f"MUTANT: growth = {mut_growth:.6f}, succ_ex = {mut_succ:.6f}")

#─── J/K) Export summary & flux comparison ──────────────────────────────

summary = {

"wild_growth": wild_growth,

"wild_succ_ex": wild_succ,

"mut_growth": mut_growth,

"mut_succ_ex": mut_succ,

"max_succ": max_succ

}

df_sum	= pd.DataFrame([summary])

df_ko	= pd.DataFrame({"knockout_rxn": ko_list})

\# flux comparison

df_flux = pd.DataFrame({

"wildtype": wild_sol.fluxes,

"mutant": mut_sol.fluxes

})

out_file = "OptKnock_iJN1463_succinate.xlsx"

with pd.ExcelWriter(out_file) as writer:

df_sum.to_excel(writer, sheet_name="Summary", index=False)

df_ko.to_excel(writer, sheet_name="Knockouts", index=False)

df_flux.to_excel(writer, sheet_name="FluxComparison")

print(f"\nResults written to {out_file}")

if __name__ == "__main__":

main()

**With RAG Code:**

import cobra

import straindesign as sd

import pandas as pd

def is_knockout_value(val):

if isinstance(val, str):

return val.strip().upper() in {"KO", "KNOCKOUT", "DEL", "DELETE"}

try:

return val < 0

except Exception:

return False

def main():

\# ----------------------

\# Load model

\# ----------------------

model = cobra.io.read_sbml_model("iJN1463.xml")

\# ----------------------

\# Set glucose uptake

\# ----------------------

if "EX_glc__D_e" in model.reactions:

model.reactions.get_by_id("EX_glc__D_e").lower_bound = -6.0

\# ----------------------

\# Set other exchanges to secretion-only

\# ----------------------

secretion_only = [

"EX_co2_e",

"EX_glcn_e",

"EX_25dkglcn_e",

"EX_catechol_e",

"EX_lac__D_e",

"EX_meoh_e",

"EX_4oxptn_e",

"EX_ac_e"

]

for rxn_id in secretion_only:

if rxn_id in model.reactions:

r = model.reactions.get_by_id(rxn_id)

r.lower_bound = 0.0

r.upper_bound = 1000.0

\# ----------------------

\# Block secretion ONLY (commented out as per your code)

\# ----------------------

block_only = []

for rxn_id in block_only:

if rxn_id in model.reactions:

r = model.reactions.get_by_id(rxn_id)

r.lower_bound = 0.0

r.upper_bound = 0.0

\# ----------------------

\# EXCLUSION LIST - prevent OptKnock from touching these

\# ----------------------

excluded_rxns = [

"ATPM",

"CAT",

"CYO1_KT",

"CYTBO3_4pp",

"CYTCAA3pp",

"NADH16pp",

"NAt3_1p5pp",

"PItex",

"PPK"

]

\# ----------------------

\# Biomass & target

\# ----------------------

biomass_rxn = "BIOMASS_KT2440_WT3"

target_rxn = "EX_succ_e"

\# verify target exists

if target_rxn not in model.reactions:

raise ValueError(f"Exchange reaction {target_rxn} not found in the model.")

\# ----------------------

\# Wild-type baseline

\# ----------------------

model.objective = biomass_rxn

wt_sol = model.optimize()

wt_flux = wt_sol.fluxes

wt_growth = wt_sol.objective_value

\# ----------------------

\# Theoretical max succinate (unconstrained)

\# ----------------------

succ_model = model.copy()

succ_model.objective = target_rxn

theoretical_succ = succ_model.optimize().objective_value

\# ----------------------

\# OptKnock module (allow up to 3 knockouts)

\# ----------------------

viability_threshold = 0.1 * wt_growth # 10% WT growth

optknock_module = sd.SDModule(

model=model,

module_type=sd.OPTKNOCK,

inner_objective=biomass_rxn,

outer_objective=target_rxn,

constraints=[f"{biomass_rxn} >= {viability_threshold}"]

)

\# ----------------------

\# Run OptKnock

\# ----------------------

max_solutions = 1

solutions = sd.compute_strain_designs(

model,

sd_modules=[optknock_module],

max_solutions=max_solutions, # number of solutions solutions

max_cost=2, # number of knockouts knockouts

solution_approach=sd.BEST

)

\# ----------------------

\# Loop over solutions, pick first one not containing excluded KOs

\# ----------------------

ko_list = []

found_valid = False

for i in range(max_solutions):

try:

reaction_sd = solutions.get_reaction_sd(i)

except Exception:

break # No more solutions

candidate_ko = []

skip = False

for rd in reaction_sd:

for rxn_id, val in rd.items():

if is_knockout_value(val):

if rxn_id in excluded_rxns:

print(f"Solution {i}: Skipped because knockout {rxn_id} is in exclusion list.")

skip = True

candidate_ko.append(rxn_id)

if skip:

continue # Try next solution

ko_list = candidate_ko

found_valid = True

print(f"Selected solution {i} with knockouts: {ko_list}")

break # Only first valid solution

if not found_valid or not ko_list:

print("No valid OptKnock solution found (all contain excluded reactions).")

return

\# ----------------------

\# Build mutant from solution

\# ----------------------

mutant = model.copy()

for rxn_id in ko_list:

if rxn_id in mutant.reactions:

mutant.reactions.get_by_id(rxn_id).knock_out()

\# ----------------------

\# Mutant simulation

\# ----------------------

mutant.objective = biomass_rxn

mut_sol = mutant.optimize()

mut_flux = mut_sol.fluxes

\# ----------------------

\# Prepare Excel output

\# ----------------------

wt_df = wt_flux.reset_index()

wt_df.columns = ["reaction_id", "wt_flux"]

mut_df = mut_flux.reset_index()

mut_df.columns = ["reaction_id", "mutant_flux"]

compare_df = wt_df.merge(mut_df, on="reaction_id", how="outer")

compare_df["delta(mut-wt)"] = compare_df["mutant_flux"] - compare_df["wt_flux"]

ko_df = pd.DataFrame({"knockout_reaction_id": ko_list})

summary_df = pd.DataFrame({

"metric": [

"WT_growth",

"WT_succinate",

"Mut_growth",

"Mut_succinate",

"theoretical_max_succinate",

"num_KOs"

],

"value": [

wt_growth,

wt_flux.get(target_rxn, float("nan")),

mut_sol.objective_value,

mut_flux.get(target_rxn, float("nan")),

theoretical_succ,

len(ko_list)

]

})

out_file = "OptKnock_iJN1463_succinate.xlsx"

with pd.ExcelWriter(out_file, engine="openpyxl") as writer:

summary_df.to_excel(writer, sheet_name="summary", index=False)

ko_df.to_excel(writer, sheet_name="knockouts", index=False)

compare_df.to_excel(writer, sheet_name="flux_comparison", index=False)

print(f"OptKnock computation complete. Results written to: {out_file}")

if __name__ == "__main__":

main()

**Task 3: Proteomics and Metabolomics Integration, Complexity: Complex**

**Protocol:**

I have uploaded the iJN1463 irreversible *Pseudomonas putida* KT2440 model (iJN1463_irrev.xml), ΔG° values (delG_post_opt.xlsx), and per-unit protein cost data (per_unit_protein_data.xlsx). Please combine ecGEM and OptMDFPathway algorithm together to ecOptMDFPathway and perform the following using your codebase and Gurobi solver:

A) Set metabolite concentration bounds: minimum 1e-6 M, maximum 100 M.

B) Set proteome capacity upper bound to 0.60 g/gDW, minimum biomass flux to 1e-6, minimum active reaction flux to 1e-6.

C) Formulate a Gurobi MILP enforcing: stoichiometric mass balance, thermodynamic driving forces from ΔG° and metabolite concentrations, flux-activity coupling via binary variables, MDF constraint (all active reactions ≥ B), proteome capacity, and minimum biomass (BIOMASS_KT2440_WT3).

D) Maximize biomass growth + weighted MDF (λ=1e-4) − weighted total flux (λ=1e-4).

E) Extract optimal fluxes, metabolite concentrations, and reaction ΔG values.

F) Compute per-reaction protein usage and total proteome requirement.

G) Export flux, concentration, thermodynamics, and protein requirement sheets to ecOptMDFPathway.xlsx.

H) Report biomass growth rate, total protein requirement, and minimum thermodynamic driving force B.

**Without RAG Code:**

#!/usr/bin/env python3

import os

import math

import cobra

import pandas as pd

import numpy as np

from gurobipy import Model, GRB, quicksum

\# Constants

R = 8.314e-3	# kJ/(mol·K)

T = 298.15	# K

RT = R * T

\# Parameters

MIN_CONC = 1e-6	# M

MAX_CONC = 100.0	# M

PROTEOME_CAPACITY = 0.60 # g protein / gDW

MIN_BIOMASS = 1e-6	# minimal biomass flux

LAM = 1e-4	# weight for MDF and total flux

\# File paths (adjust if needed)

UPLOAD_DIR	= "/app/data/uploaded_files"

MODEL_FILE	= "iJN1463_irrev.xml"

DELG_FILE	= "delG_post_opt.xlsx"

PROT_COST_FILE	= "per_unit_protein_data.xlsx"

OUTPUT_XLSX	= "ecOptMDFPathway.xlsx"

def find_file(name):

"""Helper to find uploaded file by substring."""

for root, dirs, files in os.walk(UPLOAD_DIR):

for f in files:

if name in f:

return os.path.join(root, f)

raise FileNotFoundError(f"Could not find '{name}' under {UPLOAD_DIR}")

def load_data():

"""Load ΔG° and protein cost tables."""

dg_path	= find_file("delG_post_opt.xlsx")

pc_path	= find_file("per_unit_protein_data.xlsx")

df_dg	= pd.read_excel(dg_path, index_col=0)	# ReactionID, dG0_prime (kJ/mol)

df_protc = pd.read_excel(pc_path, index_col=0) # ReactionID, kcat, MW, per‐unit protein cost (g·h/mmol)

return df_dg, df_protc

def build_milp(model, df_dg, df_protc):

"""Formulate ecOptMDFPathway MILP in Gurobi."""

\# Prepare stoichiometric matrix and index lists

mets = [m.id for m in model.metabolites]

rxns = [r.id for r in model.reactions]

S	= model.metabolites.list_attr("id")

\# Big‐M values for flux‐binary coupling

Mflux = {r.id: max(abs(r.upper_bound), abs(r.lower_bound), 1000.0) for r in model.reactions}

\# Create Gurobi model

m = Model("ecOptMDFPathway")

m.setParam('OutputFlag', 1)

\# Variables

v = m.addVars(rxns, lb={r.id: r.lower_bound for r in model.reactions},

ub={r.id: r.upper_bound for r in model.reactions},

name="v")

y = m.addVars(rxns, vtype=GRB.BINARY, name="y")

c = m.addVars(mets, lb=math.log(MIN_CONC), ub=math.log(MAX_CONC), name="log_c")

dG = m.addVars(rxns, lb=-GRB.INFINITY, ub=GRB.INFINITY, name="dG")

B = m.addVar(lb=0.0, ub=GRB.INFINITY, name="MDF")

\# 1) Stoichiometric mass balance: S · v = 0

for met in model.metabolites:

expr = quicksum(met._reaction_coefficients[rxn] * v[rxn]

for rxn in rxn for met in []) # placeholder

\# We can get stoich via model.metabolites but cobrapy does not expose directly a dict:

\# Instead do:

expr = quicksum(model.reactions.get_by_id(rxn).get_coefficient(met.id) * v[rxn] for rxn in rxns)

m.addConstr(expr == 0, name=f"massbal_{met.id}")

\# 2) Thermodynamic driving force: dG_j = dG0_j + RT * sum_i ν_ij * log(c_i)

for rid in rxns:

\# Fetch ΔG0′

dG0 = df_dg.at[rid, 'dG0_prime'] if rid in df_dg.index else 0.0

stoich = model.reactions.get_by_id(rid).metabolites # dict metabolite->stoich

therm_expr = dG0 + quicksum(RT * nu * c[met.id] for met, nu in stoich.items())

m.addConstr(dG[rid] == therm_expr, name=f"dGdef_{rid}")

\# 3) Flux‐activity coupling: v_j ≤ M_j · y_j and v_j ≥ ε · y_j

EPS = 1e-6

for rid in rxns:

m.addConstr(v[rid] <= Mflux[rid] * y[rid], name=f"coupling_pos_{rid}")

\# If reactions are irreversible and v≥0, we add minimum:

if model.reactions.get_by_id(rid).lower_bound >= 0:

m.addConstr(v[rid] >= EPS * y[rid], name=f"coupling_min_{rid}")

\# 4) MDF constraint: for all active reactions, driving force ≥ B

\#	i.e. -dG_j ≥ B · y_j → -dG_j - B·y_j ≥ 0

for rid in rxns:

m.addConstr(-dG[rid] - B * y[rid] >= 0, name=f"MDF_{rid}")

\# 5) Proteome capacity: sum_j protein_cost_j * v_j ≤ capacity

\#	protein cost (g protein · h / mmol) * mmol/gDW/h → g/gDW

prot_cost = {rid: df_protc.at[rid, 'protein_cost'] if rid in df_protc.index else 0.0 for rid in rxns}

m.addConstr(quicksum(prot_cost[rid] * v[rid] for rid in rxns) <= PROTEOME_CAPACITY,

name="proteome_capacity")

\# 6) Minimum biomass flux

BIOM = "BIOMASS_KT2440_WT3"

if BIOM not in rxns:

raise KeyError(f"Biomass reaction '{BIOM}' not found in model.")

m.addConstr(v[BIOM] >= MIN_BIOMASS, name="min_biomass")

\# Objective: maximize biomass + λ·B – λ·sum(v_j)

obj = v[BIOM] + LAM * B - LAM * quicksum(v[rid] for rid in rxns)

m.setObjective(obj, GRB.MAXIMIZE)

m.update()

return m, v, c, dG, B

def extract_results(m, v, c, dG, B):

"""Solve MILP and return results dicts."""

m.optimize()

if m.status != GRB.OPTIMAL:

raise RuntimeError("ecOptMDFPathway MILP did not find optimal solution.")

fluxes = {rid: v[rid].X for rid in v.keys()}

conc	= {mid: math.exp(c[mid].X) for mid in c.keys()}

dg	= {rid: dG[rid].X for rid in dG.keys()}

MDF	= B.X

return fluxes, conc, dg, MDF

def compute_protein_usage(fluxes, df_protc):

"""Compute per‐reaction protein usage and sum total."""

usage = {}

for rid, v in fluxes.items():

if rid in df_protc.index:

cost = df_protc.at[rid, 'protein_cost'] # g·h/mmol

usage[rid] = cost * max(v, 0)

else:

usage[rid] = 0.0

total = sum(usage.values())

return usage, total

def write_excel(fluxes, conc, dg, usage, total_prot, MDF, growth):

"""Save multiple sheets into one Excel workbook."""

with pd.ExcelWriter(OUTPUT_XLSX) as writer:

pd.DataFrame.from_dict(fluxes, orient='index', columns=['flux']).to_excel(writer, sheet_name="Fluxes")

pd.DataFrame.from_dict(conc, orient='index', columns=['concentration_M']).to_excel(writer, sheet_name="Concentrations")

pd.DataFrame.from_dict(dg, orient='index', columns=['dG_kJ_per_mol']).to_excel(writer, sheet_name="Thermodynamics")

pd.DataFrame.from_dict(usage, orient='index', columns=['protein_g_per_gDW']).to_excel(writer, sheet_name="ProteinUsage")

\# Summary sheet

summary = {

'biomass_h-1': growth,

'total_protein_g/gDW': total_prot,

'MDF_kJ_per_mol': MDF

}

pd.DataFrame([summary]).to_excel(writer, sheet_name="Summary", index=False)

print(f"Results written to {OUTPUT_XLSX}")

def main():

\# 1) Load SBML and data tables

model_path = find_file("iJN1463_irrev.xml")

model = cobra.io.read_sbml_model(model_path)

df_dg, df_protc = load_data()

\# 2) Build and solve MILP

milp, v_vars, c_vars, dG_vars, B_var = build_milp(model, df_dg, df_protc)

fluxes, conc, dg_vals, MDF = extract_results(milp, v_vars, c_vars, dG_vars, B_var)

\# 3) Extract biomass growth

growth = fluxes["BIOMASS_KT2440_WT3"]

\# 4) Compute protein usage

usage, total_prot = compute_protein_usage(fluxes, df_protc)

\# 5) Write Excel output

write_excel(fluxes, conc, dg_vals, usage, total_prot, MDF, growth)

if __name__ == "__main__":

main()

**With RAG Code:**

import cobra

import pandas as pd

import numpy as np

import gurobipy as gp

from gurobipy import GRB

\# ============================================================

\# PARAMETERS

\# ============================================================

PARAMS = {

"MODEL_SBML": "iJN1463_irrev.xml",

"DELG_XLSX": "delG_post_opt.xlsx",

"PROTEIN_XLSX": "per_unit_protein_data.xlsx",

"BIOMASS_RXN": "BIOMASS_KT2440_WT3",

"OUTPUT_XLSX": "ecOptMDFPathway.xlsx",

"R_kJ_per_mol_K": 8.314e-3,

"T_K": 298.15,

"Cmin": 1e-6,

"Cmax": 100.0,

"PROTEOME_UB_g_per_gDW": 0.60,

"PROTEIN_COST_DIVISOR": 1000.0,

"lambda_B": 1e-4,

"lambda_F": 1e-4,

"min_biomass": 1e-6,

"min_flux": 1e-6,

"BIG_M": 1000,

"gurobi_OutputFlag": 0,

"gurobi_DualReductions": 0,

}

\# ============================================================

\# Gibbs calculation

\# ============================================================

def compute_delG_from_solution(model, delG0_dict, lnC_vals, RT):

delG = {}

for rxn in model.reactions:

s = 0.0

for met, nu in rxn.metabolites.items():

s += float(nu) * float(lnC_vals[met.id])

delG[rxn.id] = float(delG0_dict[rxn.id]) + RT * s

return delG

\# ============================================================

\# MODEL BUILD

\# ============================================================

def build_and_solve(model, delG0, prot_cost, params):

Rg = params["R_kJ_per_mol_K"]

T = params["T_K"]

RT = Rg * T

x_lb = np.log(params["Cmin"])

x_ub = np.log(params["Cmax"])

BIG_M = params["BIG_M"]

m = gp.Model("ecOptMDFPathway")

m.setParam("OutputFlag", params["gurobi_OutputFlag"])

m.setParam("DualReductions", params["gurobi_DualReductions"])

\# --------------------------------------------------------

\# VARIABLES

\# --------------------------------------------------------

v = {}

y = {}

for rxn in model.reactions:

ub = max(0.0, float(rxn.upper_bound))

v[rxn.id] = m.addVar(lb=0, ub=ub, name=f"v_{rxn.id}")

y[rxn.id] = m.addVar(vtype=GRB.BINARY, name=f"y_{rxn.id}")

x = {met.id: m.addVar(lb=x_lb, ub=x_ub, name=f"x_{met.id}") for met in model.metabolites}

f = {rxn.id: m.addVar(lb=-GRB.INFINITY, name=f"f_{rxn.id}") for rxn in model.reactions}

B = m.addVar(lb=0, name="B")

m.update()

\# --------------------------------------------------------

\# MASS BALANCE

\# --------------------------------------------------------

for met in model.metabolites:

m.addConstr(

gp.quicksum(float(rxn.metabolites[met]) * v[rxn.id] for rxn in met.reactions) == 0,

name=f"mass_{met.id}",

)

\# --------------------------------------------------------

\# THERMODYNAMICS

\# --------------------------------------------------------

for rxn in model.reactions:

dG0 = float(delG0[rxn.id])

m.addConstr(

f[rxn.id] == -(dG0 + RT * gp.quicksum(float(coeff) * x[met.id] for met, coeff in rxn.metabolites.items())),

name=f"thermo_{rxn.id}",

)

\# --------------------------------------------------------

\# FLUX-ACTIVITY COUPLING

\# --------------------------------------------------------

for rxn in model.reactions:

ub = max(0.0, float(rxn.upper_bound))

m.addConstr(v[rxn.id] <= ub * y[rxn.id], name=f"flux_ub_{rxn.id}")

m.addConstr(v[rxn.id] >= params["min_flux"] * y[rxn.id], name=f"flux_lb_{rxn.id}")

\# --------------------------------------------------------

\# MDF COUPLING

\# --------------------------------------------------------

for rxn in model.reactions:

m.addConstr(f[rxn.id] >= B - BIG_M * (1 - y[rxn.id]), name=f"mdf_{rxn.id}")

\# --------------------------------------------------------

\# PROTEOME CONSTRAINT

\# --------------------------------------------------------

if prot_cost:

prot_expr = gp.quicksum(float(cost) * v[rid] for rid, cost in prot_cost.items()) / params["PROTEIN_COST_DIVISOR"]

m.addConstr(prot_expr <= params["PROTEOME_UB_g_per_gDW"], name="protein_capacity")

\# --------------------------------------------------------

\# MINIMUM BIOMASS

\# --------------------------------------------------------

biomass_id = params["BIOMASS_RXN"]

if params["min_biomass"] > 0:

m.addConstr(v[biomass_id] >= params["min_biomass"], name="min_biomass")

\# --------------------------------------------------------

\# OBJECTIVE

\# --------------------------------------------------------

total_flux = gp.quicksum(v[rxn.id] for rxn in model.reactions)

m.setObjective(v[biomass_id] + params["lambda_B"] * B - params["lambda_F"] * total_flux, GRB.MAXIMIZE)

m.optimize()

return m, v, x, B, RT

\# ============================================================

\# LOAD MODEL

\# ============================================================

model = cobra.io.read_sbml_model(PARAMS["MODEL_SBML"])

if PARAMS["BIOMASS_RXN"] not in [r.id for r in model.reactions]:

raise ValueError("Biomass reaction not found.")

\# ============================================================

\# LOAD ΔG°

\# ============================================================

delG_df = pd.read_excel(PARAMS["DELG_XLSX"])

delG0 = pd.Series(delG_df["delG0"].values, index=delG_df["Reaction"]).to_dict()

\# ============================================================

\# LOAD PROTEIN COST

\# ============================================================

prot_raw = pd.read_excel(PARAMS["PROTEIN_XLSX"], usecols=[0, 1], names=["Reactions", "Protein_cost"])

model_rxns = set(r.id for r in model.reactions)

prot_cost = {}

for _, row in prot_raw.iterrows():

rid = str(row["Reactions"])

if rid in model_rxns and pd.notna(row["Protein_cost"]):

prot_cost[rid] = float(row["Protein_cost"])

\# ============================================================

\# SOLVE

\# ============================================================

m, v, x, B, RT = build_and_solve(model, delG0, prot_cost, PARAMS)

if m.status != GRB.OPTIMAL:

raise RuntimeError("Optimization did not converge.")

\# ============================================================

\# EXTRACT SOLUTION

\# ============================================================

fluxes = {rid: var.X for rid, var in v.items()}

lnC = {mid: var.X for mid, var in x.items()}

C = {mid: np.exp(val) for mid, val in lnC.items()}

delG_model = compute_delG_from_solution(model, delG0, lnC, RT)

\# ============================================================

\# DATAFRAMES

\# ============================================================

flux_df = pd.DataFrame.from_dict(fluxes, orient="index", columns=["flux"]).reset_index()

flux_df.columns = ["Reaction", "flux"]

conc_df = pd.DataFrame.from_dict(C, orient="index", columns=["concentration"]).reset_index()

conc_df.columns = ["Metabolite", "concentration"]

thermo_df = pd.DataFrame({

"Reaction": [r.id for r in model.reactions],

"delG0_final": [delG0[r.id] for r in model.reactions],

"delG_model": [delG_model[r.id] for r in model.reactions],

})

\# ============================================================

\# PROTEIN USAGE

\# ============================================================

protein_rows = []

p_total = 0.0

for rid, cost in prot_cost.items():

pr = cost * fluxes.get(rid, 0) / PARAMS["PROTEIN_COST_DIVISOR"]

protein_rows.append({

"Reaction": rid,

"Protein_Required (g/gDW)": pr

})

p_total += pr

protein_df = pd.DataFrame(protein_rows)

\# ============================================================

\# WRITE EXCEL

\# ============================================================

with pd.ExcelWriter(PARAMS["OUTPUT_XLSX"], engine="openpyxl") as writer:

flux_df.to_excel(writer, sheet_name="flux", index=False)

conc_df.to_excel(writer, sheet_name="concentration", index=False)

thermo_df.to_excel(writer, sheet_name="thermodynamics", index=False)

protein_df.to_excel(writer, sheet_name="protein_requirements", index=False)

print("Biomass growth rate (/hr) =", v[PARAMS["BIOMASS_RXN"]].X)

print(f"Total protein requirement (g/gDW): {p_total}")

print("Minimum thermodynamic driving force (Kj/mol) =", B.X)

**ChatGEM Evaluation Report by Claude Sonnet 4.6: RAG vs. Without RAG**

Evaluated by Claude Sonnet 4.6

Overall Performance Score (OPS) = 0.7 x Coding Accuracy + 0.3 x Code Completeness

**1. Scoring Summary**

| **Task** | **Complexity** | **Condition** | **Accuracy (1-5)** | **Completeness (1-5)** | **OPS** |
| --- | --- | --- | --- | --- | --- |
| Task 1: FBA and Knockout | Low | Without RAG | 3.00 | 5.00 | 3.60 |
|  |  | With RAG | 4.00 | 5.00 | 4.30 |
| Task 2: OptKnock Strain Design | High | Without RAG | 2.00 | 3.00 | 2.30 |
|  |  | With RAG | 4.00 | 4.00 | 4.00 |
| Task 3: ecOptMDFPathway | Complex | Without RAG | 2.00 | 2.00 | 2.00 |
|  |  | With RAG | 4.00 | 5.00 | 4.30 |

**Scoring Rubric**

Coding Accuracy (1-5): 5 = runs correctly with expected output and follows best practices; 4 = runs correctly but with minor code quality or convention issues; 3 = runs but produces incorrect output; 2 = partially executes or uses wrong library; 1 = does not run.

Code Completeness (1-5): 5 = all required steps present; 4 = minor step or output omission; 3 = key step missing; 2 = major gaps; 1 = skeleton only.

**2. Findings by Task**

**Task 1: FBA and Knockout**

**Without RAG:** Used regex-based succinate detection instead of the exact reaction ID (EX_succ_e). Modified the model in-place without a context manager. Applied bounds=(0,0) directly instead of knock_out(). Functionally produces output but is fragile and not COBRApy best practice.

**With RAG:** Verified functional. Uses exact EX_succ_e ID and knock_out() correctly. However, the model is modified in-place without a context manager, meaning knockouts are permanent on the model object. This is a minor code quality issue that does not affect correctness for this isolated task but would cause errors in a pipeline context.

**Task 2: OptKnock Strain Design**

**Without RAG:** Used Cameo's OptKnock instead of the specified StrainDesign toolbox, directly violating the protocol. All logical steps are structurally present but the wrong library renders the code non-compliant.

**With RAG:** Verified functional. Uses the correct StrainDesign toolbox with proper sd.OPTKNOCK formulation. Minor issues: the exclusion list logic appends rxn_id to candidate_ko before evaluating the skip flag, meaning excluded reactions are still recorded in the candidate list even when the solution is skipped. Additionally, the flux comparison sheet lacks a delta column as implied by the protocol.

**Task 3: ecOptMDFPathway**

**Without RAG:** Contains a critical bug in the mass balance loop (broken list comprehension placeholder that is immediately overwritten). MDF constraint formulation is incorrect (-dG - B*y >= 0 does not correctly enforce the Big-M relaxation). Column names are assumed and may not match actual data files. Code will not execute as written.

**With RAG:** Verified functional. Clean Gurobi MILP with correct mass balance, flux-activity coupling, proteome constraint, and objective. Minor issue: the thermodynamic driving force variable f is defined as the negation of the standard dG expression, which is internally consistent but deviates from conventional sign convention, reducing interpretability. All output sheets are exported correctly.

**3. Overall Comparison**

| **Metric** | **Without RAG** | **With RAG** |
| --- | --- | --- |
| Mean Accuracy | 2.33 | 4.00 |
| Mean Completeness | 3.33 | 4.67 |
| Mean OPS | 2.63 | 4.20 |
