## Supplementary material for "ChatGEM: An Agentic Architecture Enabling Interactive Simulation of Genome-Scale Metabolic Models": To evaluate whether model choice affects code generation performance, we compared the default o4-mini, a lightweight reasoning model, against GPT-4.1, (from another version of this preprint: version 1, 2026-07-21)

**Model: ChatGPT 4.1**

**Task1: FBA and knockout; Complexity: Low**

**Protocol:**

I have uploaded the iJN1463.xml model of *Pseudomonas putida* KT2440. Please perform the following using the codebase and Gurobi solver:

A) Run baseline FBA and record biomass growth rate and succinate exchange flux.

B) Knock out reactions: FUM, ATPS4rpp, HCO3E, H2CO3D.

C) Rerun FBA after knockouts.

D) Report biomass and succinate exchange flux before and after deletions

**Without RAG Code:**

import cobra

\# Load the model

model = cobra.io.read_sbml_model("iJN1463.xml")

\# Set Gurobi as the solver

model.solver = "gurobi"

\# --- Helper function ---

def run_fba_and_report(model, label):

solution = model.optimize()

\# Adjust these IDs if different in your SBML:

biomass_rxn = model.reactions.get_by_id("BIOMASS")

succ_ex_rxn = model.reactions.get_by_id("EX_succ_e") # or check your actual exchange reaction

print(f"Results [{label}]:")

print(f" Biomass growth rate: {solution.objective_value:.4f}")

print(f" Succinate exchange flux: {succ_ex_rxn.flux:.4f}\n")

return solution.objective_value, succ_ex_rxn.flux

\# ---- (A) Baseline ----

print("A) Baseline FBA:")

biomass_before, succ_flux_before = run_fba_and_report(model, "Baseline")

\# ---- (B) Knockouts ----

\# List of reaction IDs to knock out

ko_list = ["FUM", "ATPS4rpp", "HCO3E", "H2CO3D"]

print("B) Deleting reactions:", ko_list)

for rxn_id in ko_list:

model.reactions.get_by_id(rxn_id).knock_out()

\# ---- (C)+(D) FBA after knockouts + report ----

print("C+D) FBA after knockouts:")

biomass_after, succ_flux_after = run_fba_and_report(model, "KO")

print("Summary Table:")

print(f"{'Condition':<12} {'Biomass':<10} {'Succ. exchange':<16}")

print(f"{'Baseline':<12} {biomass_before:<10.4f} {succ_flux_before:<16.4f}")

print(f"{'After KO':<12} {biomass_after:<10.4f} {succ_flux_after:<16.4f}")

**With RAG Code:**

from cobra.io import read_sbml_model

model = read_sbml_model("iJN1463.xml")

model.solver = "gurobi"

sol = model.optimize()

print("Before")

print("Biomass =", sol.objective_value)

print("Succinate =", sol.fluxes["EX_succ_e"])

for rxn in ["FUM", "ATPS4rpp", "HCO3E", "H2CO3D"]:

model.reactions.get_by_id(rxn).knock_out()

sol = model.optimize()

print("\nAfter")

print("Biomass =", sol.objective_value)

print("Succinate =", sol.fluxes["EX_succ_e"])

**Task 2: Strain design by OptKnock, Complexity: High**

**Protocol:**

I have uploaded the iJN1463.xml model of *Pseudomonas putida* KT2440. Please perform the following task using the codebase with StrainDesign toolbox and Gurobi solver:

A) Set glucose uptake to -6.0 mmol/gDW/h.

B) Set the following exchanges to secretion-only (lower bound = 0): EX_co2_e, EX_glcn_e, EX_25dkglcn_e, EX_catechol_e, EX_lac__D_e, EX_meoh_e, EX_4oxptn_e, EX_ac_e.

C) Set biomass objective to BIOMASS_KT2440_WT3 and target to EX_succ_e.

D) Run baseline FBA and record wild-type biomass growth rate and succinate exchange flux.

E) Calculate theoretical maximum succinate by optimizing EX_succ_e directly.

F) Run OptKnock with maximum 2 knockouts and 1 solution, maximizing succinate (outer objective) while maintaining at least 10% wild-type growth (inner objective).

G) Exclude reactions ATPM, CAT, CYO1_KT, CYTBO3_4pp, CYTCAA3pp, NADH16pp, NAt3_1p5pp, PItex, PPK from knockout candidates.

H) Select the first valid solution not containing excluded reactions.

I) Apply knockouts to a mutant model and simulate growth.

J) Report wild-type vs mutant biomass and succinate flux, knockout reactions, and theoretical maximum succinate.

K) Export summary, knockout list, and full flux comparison to OptKnock_iJN1463_succinate.xlsx.

**Without RAG Code:**

from strain_design.model import CobraModel

from strain_design.strain_optimization import OptKnock

from strain_design.export import export_flux_comparison

from strain_design.simulation import set_medium_exchanges

import pandas as pd

\# (A) Load iJN1463 model and set solver

model = CobraModel.from_sbml("iJN1463.xml")

model.solver = "gurobi"

\# (A) Set glucose uptake to -6 mmol/gDW/h

glc = model.reactions.get_by_id("EX_glc__D_e")

glc.lower_bound = -6.0

glc.upper_bound = 0.0 # If you want to make it uptake only

\# (B) Set selected exchanges to secretion-only

secretion_only = [

"EX_co2_e", "EX_glcn_e", "EX_25dkglcn_e", "EX_catechol_e",

"EX_lac__D_e", "EX_meoh_e", "EX_4oxptn_e", "EX_ac_e"

]

for rxn_id in secretion_only:

rxn = model.reactions.get_by_id(rxn_id)

rxn.lower_bound = 0.0

\# (C) Set objectives and target

model.objective = "BIOMASS_KT2440_WT3"

target_rxn_id = "EX_succ_e"

\# (D) Baseline FBA

sol_wt = model.optimize()

wt_biomass = sol_wt.objective_value

wt_succ_flux = model.reactions.get_by_id(target_rxn_id).flux

print(f"Wild-type: Biomass = {wt_biomass:.3f}	Succinate = {wt_succ_flux:.3f}")

\# (E) Maximize succinate

model.objective = target_rxn_id

sol_succ = model.optimize()

theoretical_succ = sol_succ.objective_value

print(f"Theoretical max succinate = {theoretical_succ:.3f}")

\# (F) OptKnock setup: max 2 KOs, maintain >=10% WT growth

ok = OptKnock(model)

excluded = {

"ATPM","CAT","CYO1_KT","CYTBO3_4pp","CYTCAA3pp","NADH16pp","NAt3_1p5pp","PItex","PPK"

}

knockout_candidates = [

rxn.id for rxn in model.reactions

if rxn.id not in excluded and not rxn.reversibility is False # Exclude irreversible importers

]

ok.target = target_rxn_id

ok.max_knockouts = 2

ok.inner_objective = "BIOMASS_KT2440_WT3"

ok.constraints = [{ "BIOMASS_KT2440_WT3": (0.1 * wt_biomass, None) }]

ok.candidates = knockout_candidates

ok.timelimit = 600 # Optional: set a time limit

\# (F) Run OptKnock

solutions = ok.run(max_solutions=1)

ko_sol = solutions[0]

\# (H) If solution includes only valid KOs

if all(rxn not in excluded for rxn in ko_sol["knockouts"]):

knockouts = list(ko_sol["knockouts"])

else:

knockouts = [rxn for rxn in ko_sol["knockouts"] if rxn not in excluded] # Conservative fallback

print(f"OptKnock knockouts: {knockouts}")

\# (I) Apply knockouts to mutant

mutant = model.copy() # Use a fresh copy of the WT model for mutant

mutant.objective = "BIOMASS_KT2440_WT3"

for rxn_id in knockouts:

mutant.reactions.get_by_id(rxn_id).knock_out()

sol_mut = mutant.optimize()

mut_biomass = sol_mut.objective_value

mut_succ_flux = mutant.reactions.get_by_id(target_rxn_id).flux

\# (J) Print summary

print("--- SUMMARY ---")

print(f"Wild-type biomass: {wt_biomass:.3f}")

print(f"Wild-type succinate: {wt_succ_flux:.3f}")

print(f"Mutant biomass: {mut_biomass:.3f}")

print(f"Mutant succinate: {mut_succ_flux:.3f}")

print(f"Theoretical max succinate: {theoretical_succ:.3f}")

print(f"Knockout RXNs: {knockouts}")

\# (K) Export results to Excel

\# Flux comparison (WT vs KO)

wt_flux = pd.DataFrame({"WT": sol_wt.fluxes})

mut_flux = pd.DataFrame({"Mutant": sol_mut.fluxes})

all_flux = pd.concat([wt_flux, mut_flux], axis=1)

all_flux.index.name = "Reaction"

\# Write summary and lists

with pd.ExcelWriter("OptKnock_iJN1463_succinate.xlsx") as writer:

all_flux.to_excel(writer, sheet_name="Fluxes")

\# Summary Table

summary = pd.DataFrame({

"Condition": ["Wild-type", "Mutant", "Max-succinate"],

"Biomass": [wt_biomass, mut_biomass, None],

"Succinate": [wt_succ_flux, mut_succ_flux, theoretical_succ]

})

summary.to_excel(writer, sheet_name="Summary", index=False)

\# Knockouts

pd.DataFrame(knockouts, columns=["Knockout reactions"]).to_excel(writer, sheet_name="Knockouts", index=False)

print("Results exported to OptKnock_iJN1463_succinate.xlsx")

**With RAG Code:**

import cobra

import straindesign as sd

import pandas as pd

def is_knockout_value(val):

if isinstance(val, str):

return val.strip().upper() in {"KO", "KNOCKOUT", "DEL", "DELETE"}

try:

return val < 0

except Exception:

return False

def main():

\# ----------------------

\# Load model

\# ----------------------

model = cobra.io.read_sbml_model("iJN1463.xml")

\# ----------------------

\# Set glucose uptake

\# ----------------------

if "EX_glc__D_e" in model.reactions:

model.reactions.get_by_id("EX_glc__D_e").lower_bound = -6.0

\# ----------------------

\# Set other exchanges to secretion-only

\# ----------------------

secretion_only = [

"EX_co2_e",

"EX_glcn_e",

"EX_25dkglcn_e",

"EX_catechol_e",

"EX_lac__D_e",

"EX_meoh_e",

"EX_4oxptn_e",

"EX_ac_e"

]

for rxn_id in secretion_only:

if rxn_id in model.reactions:

r = model.reactions.get_by_id(rxn_id)

r.lower_bound = 0.0

r.upper_bound = 1000.0

\# ----------------------

\# Block secretion ONLY (commented out as per your code)

\# ----------------------

block_only = []

for rxn_id in block_only:

if rxn_id in model.reactions:

r = model.reactions.get_by_id(rxn_id)

r.lower_bound = 0.0

r.upper_bound = 0.0

\# ----------------------

\# EXCLUSION LIST - prevent OptKnock from touching these

\# ----------------------

excluded_rxns = [

"ATPM",

"CAT",

"CYO1_KT",

"CYTBO3_4pp",

"CYTCAA3pp",

"NADH16pp",

"NAt3_1p5pp",

"PItex",

"PPK"

]

\# ----------------------

\# Biomass & target

\# ----------------------

biomass_rxn = "BIOMASS_KT2440_WT3"

target_rxn = "EX_succ_e"

\# verify target exists

if target_rxn not in model.reactions:

raise ValueError(f"Exchange reaction {target_rxn} not found in the model.")

\# ----------------------

\# Wild-type baseline

\# ----------------------

model.objective = biomass_rxn

wt_sol = model.optimize()

wt_flux = wt_sol.fluxes

wt_growth = wt_sol.objective_value

\# ----------------------

\# Theoretical max succinate (unconstrained)

\# ----------------------

succ_model = model.copy()

succ_model.objective = target_rxn

theoretical_succ = succ_model.optimize().objective_value

\# ----------------------

\# OptKnock module (allow up to 3 knockouts)

\# ----------------------

viability_threshold = 0.1 * wt_growth # 10% WT growth

optknock_module = sd.SDModule(

model=model,

module_type=sd.OPTKNOCK,

inner_objective=biomass_rxn,

outer_objective=target_rxn,

constraints=[f"{biomass_rxn} >= {viability_threshold}"]

)

\# ----------------------

\# Run OptKnock

\# ----------------------

max_solutions = 1

solutions = sd.compute_strain_designs(

model,

sd_modules=[optknock_module],

max_solutions=max_solutions, # number of solutions solutions

max_cost=2, # number of knockouts knockouts

solution_approach=sd.BEST

)

\# ----------------------

\# Loop over solutions, pick first one not containing excluded KOs

\# ----------------------

ko_list = []

found_valid = False

for i in range(max_solutions):

try:

reaction_sd = solutions.get_reaction_sd(i)

except Exception:

break # No more solutions

candidate_ko = []

skip = False

for rd in reaction_sd:

for rxn_id, val in rd.items():

if is_knockout_value(val):

if rxn_id in excluded_rxns:

print(f"Solution {i}: Skipped because knockout {rxn_id} is in exclusion list.")

skip = True

candidate_ko.append(rxn_id)

if skip:

continue # Try next solution

ko_list = candidate_ko

found_valid = True

print(f"Selected solution {i} with knockouts: {ko_list}")

break # Only first valid solution

if not found_valid or not ko_list:

print("No valid OptKnock solution found (all contain excluded reactions).")

return

\# ----------------------

\# Build mutant from solution

\# ----------------------

mutant = model.copy()

for rxn_id in ko_list:

if rxn_id in mutant.reactions:

mutant.reactions.get_by_id(rxn_id).knock_out()

\# ----------------------

\# Mutant simulation

\# ----------------------

mutant.objective = biomass_rxn

mut_sol = mutant.optimize()

mut_flux = mut_sol.fluxes

\# ----------------------

\# Prepare Excel output

\# ----------------------

wt_df = wt_flux.reset_index()

wt_df.columns = ["reaction_id", "wt_flux"]

mut_df = mut_flux.reset_index()

mut_df.columns = ["reaction_id", "mutant_flux"]

compare_df = wt_df.merge(mut_df, on="reaction_id", how="outer")

compare_df["delta(mut-wt)"] = compare_df["mutant_flux"] - compare_df["wt_flux"]

ko_df = pd.DataFrame({"knockout_reaction_id": ko_list})

summary_df = pd.DataFrame({

"metric": [

"WT_growth",

"WT_succinate",

"Mut_growth",

"Mut_succinate",

"theoretical_max_succinate",

"num_KOs"

],

"value": [

wt_growth,

wt_flux.get(target_rxn, float("nan")),

mut_sol.objective_value,

mut_flux.get(target_rxn, float("nan")),

theoretical_succ,

len(ko_list)

]

})

out_file = "OptKnock_iJN1463_succinate.xlsx"

with pd.ExcelWriter(out_file, engine="openpyxl") as writer:

summary_df.to_excel(writer, sheet_name="summary", index=False)

ko_df.to_excel(writer, sheet_name="knockouts", index=False)

compare_df.to_excel(writer, sheet_name="flux_comparison", index=False)

print(f"OptKnock computation complete. Results written to: {out_file}")

if __name__ == "__main__":

main()

**Task 3: Proteomics and Metabolomics Integration, Complexity: Complex**

**Protocol:**

I have uploaded the iJN1463 irreversible *Pseudomonas putida* KT2440 model (iJN1463_irrev.xml), ΔG° values (delG_post_opt.xlsx), and per-unit protein cost data (per_unit_protein_data.xlsx). Please combine ecGEM and OptMDFPathway algorithm together to ecOptMDFPathway and perform the following using your codebase and Gurobi solver:

A) Set metabolite concentration bounds: minimum 1e-6 M, maximum 100 M.

B) Set proteome capacity upper bound to 0.60 g/gDW, minimum biomass flux to 1e-6, minimum active reaction flux to 1e-6.

C) Formulate a Gurobi MILP enforcing: stoichiometric mass balance, thermodynamic driving forces from ΔG° and metabolite concentrations, flux-activity coupling via binary variables, MDF constraint (all active reactions ≥ B), proteome capacity, and minimum biomass (BIOMASS_KT2440_WT3).

D) Maximize biomass growth + weighted MDF (λ=1e-4) − weighted total flux (λ=1e-4).

E) Extract optimal fluxes, metabolite concentrations, and reaction ΔG values.

F) Compute per-reaction protein usage and total proteome requirement.

G) Export flux, concentration, thermodynamics, and protein requirement sheets to ecOptMDFPathway.xlsx.

H) Report biomass growth rate, total protein requirement, and minimum thermodynamic driving force B.

**Without RAG Code:**

import pandas as pd

import cobra

from gurobipy import Model, GRB, quicksum

import numpy as np

\# --- A) Load model and data ---

model = cobra.io.read_sbml_model("iJN1463_irrev.xml")

dG0_df = pd.read_excel("delG_post_opt.xlsx") # Columns: Reaction, dG0

prot_df = pd.read_excel("per_unit_protein_data.xlsx") # Columns: Reaction, protein_cost

reaction_ids = [r.id for r in model.reactions]

metab_ids	= [m.id for m in model.metabolites]

dG0_dict	= dict(zip(dG0_df["Reaction"], dG0_df["dG0"]))

prot_cost	= dict(zip(prot_df["Reaction"], prot_df["protein_cost"]))

\# --- A) Metabolite concentration bounds ---

conc_min = 1e-6

conc_max = 100.0

\# --- B) Proteome and flux lower bounds ---

prot_cap	= 0.60	# g protein / gDW

min_biomass = 1e-6

min_flux	= 1e-6

\# --- C) Setup Gurobi MILP ---

m = Model("ecOptMDFPathway")

m.setParam('OutputFlag', 1)

m.setParam('MIPGap', 1e-4) # Tolerance

\# --- Variables ---

\# Fluxes (v), log-conc (lc), binary activity indicator (y), single MDF variable (B)

v = m.addVars(reaction_ids, lb=-GRB.INFINITY, ub=GRB.INFINITY, name="v")

lc = m.addVars(metab_ids, lb=np.log(conc_min), ub=np.log(conc_max), name="log_c")

y = m.addVars(reaction_ids, vtype=GRB.BINARY, name="y")

dG = m.addVars(reaction_ids, lb=-GRB.INFINITY, ub=GRB.INFINITY, name="dG")

B = m.addVar(lb=0, ub=1000, name="B")

\# --- Stoichiometric Mass Balance ---

S = cobra.util.create_stoichiometric_matrix(model)

for i, met in enumerate(metab_ids):

stoich = {r.id: S[i, j] for j, r in enumerate(model.reactions) if abs(S[i, j]) > 1e-12}

m.addConstr(quicksum(v[rxn_id]*stoich[rxn_id] for rxn_id in stoich) == 0, "massbal_" + met)

\# --- Flux/Activity coupling ---

M = 1000 # Big-M

for rxn in reaction_ids:

lb = model.reactions.get_by_id(rxn).lower_bound

ub = model.reactions.get_by_id(rxn).upper_bound

\# Couple flux and activity

m.addConstr(v[rxn] >= min_flux * y[rxn], f"minFlux_{rxn}")

if ub < GRB.INFINITY:

m.addConstr(v[rxn] <= ub * y[rxn], f"ubFlux_{rxn}")

if lb > -GRB.INFINITY and lb < 0:

m.addConstr(v[rxn] >= lb * y[rxn], f"lbFlux_{rxn}") # For irreversible negative fluxes too

\# --- Biomass and minimums ---

biomass_rxn = "BIOMASS_KT2440_WT3"

m.addConstr(v[biomass_rxn] >= min_biomass, "min_biomass")

\# --- Thermodynamic constraints ---

R = 8.31e-3 # kJ/(mmol*K)

T = 298.15	# K

for rxn in reaction_ids:

rxn_obj = model.reactions.get_by_id(rxn)

if rxn in dG0_dict:

dG0 = dG0_dict[rxn]

\# dG = dG0 + RT logQ, logQ = sum(c_i^Sij)

stoich = rxn_obj.metabolites

m.addConstr(

dG[rxn] == dG0 + R*T * quicksum(Sij * lc[met.id] for met, Sij in stoich.items()), f"dGcalc_{rxn}"

)

\# dG >= B for all active reactions

m.addConstr(dG[rxn] >= B - M*(1 - y[rxn]), f"MDF_{rxn}") # If y=1 (active), dG >= B

\# --- Proteome constraint ---

total_prot = quicksum(

abs(v[rxn]) * prot_cost[rxn] if rxn in prot_cost else 0 for rxn in reaction_ids

)

m.addConstr(total_prot <= prot_cap, "proteome_constraint")

\# --- Objective ---

lam_mdf = 1e-4

lam_flux = 1e-4

obj = v[biomass_rxn] + lam_mdf * B - lam_flux * quicksum(abs(v[rxn]) for rxn in reaction_ids)

m.setObjective(obj, GRB.MAXIMIZE)

\# --- Solve MILP ---

m.optimize()

\# --- E,F) Extract result ---

if m.status == GRB.OPTIMAL:

flux_sol = {rxn: v[rxn].X for rxn in reaction_ids}

logc_sol = {m_id: lc[m_id].X for m_id in metab_ids}

conc_sol = {m_id: np.exp(lc[m_id].X) for m_id in metab_ids}

dG_sol	= {rxn: dG[rxn].X if rxn in dG else None for rxn in reaction_ids}

prot_sol = {rxn: abs(flux_sol[rxn]) * prot_cost.get(rxn, 0.) for rxn in reaction_ids}

total_prot_used = sum(prot_sol.values())

B_val	= B.X

biomass	= flux_sol[biomass_rxn]

\# --- G) Export results ---

flux_df = pd.DataFrame(list(flux_sol.items()), columns=["Reaction", "Flux"])

conc_df = pd.DataFrame(list(conc_sol.items()), columns=["Metabolite", "Concentration"])

dG_df	= pd.DataFrame([(k, dG_sol[k]) for k in dG_sol], columns=["Reaction", "dG"])

prot_df = pd.DataFrame(list(prot_sol.items()), columns=["Reaction", "ProteinUsage"])

with pd.ExcelWriter("ecOptMDFPathway.xlsx") as writer:

flux_df.to_excel(writer, sheet_name="Fluxes", index=False)

conc_df.to_excel(writer, sheet_name="Concentrations", index=False)

dG_df.to_excel(writer, sheet_name="Thermodynamics", index=False)

prot_df.to_excel(writer, sheet_name="ProteinRequirements", index=False)

\# --- H) Report ---

print(f"Optimal biomass growth rate: {biomass:.5f} 1/h")

print(f"Total protein requirement: {total_prot_used:.4f} g/gDW")

print(f"Minimum thermodynamic driving force (MDF) B: {B_val:.4f} kJ/mol")

else:

print("No optimal solution found.")

**With RAG Code:**

import cobra

import pandas as pd

import numpy as np

import gurobipy as gp

from gurobipy import GRB

\# ============================================================

\# PARAMETERS

\# ============================================================

PARAMS = {

"MODEL_SBML": "iJN1463_irrev.xml",

"DELG_XLSX": "delG_post_opt.xlsx",

"PROTEIN_XLSX": "per_unit_protein_data.xlsx",

"BIOMASS_RXN": "BIOMASS_KT2440_WT3",

"OUTPUT_XLSX": "ecOptMDFPathway.xlsx",

"R_kJ_per_mol_K": 8.314e-3,

"T_K": 298.15,

"Cmin": 1e-6,

"Cmax": 100.0,

"PROTEOME_UB_g_per_gDW": 0.60,

"PROTEIN_COST_DIVISOR": 1000.0,

"lambda_B": 1e-4,

"lambda_F": 1e-4,

"min_biomass": 1e-6,

"min_flux": 1e-6,

"BIG_M": 1000,

"gurobi_OutputFlag": 0,

"gurobi_DualReductions": 0,

}

\# ============================================================

\# Gibbs calculation

\# ============================================================

def compute_delG_from_solution(model, delG0_dict, lnC_vals, RT):

delG = {}

for rxn in model.reactions:

s = 0.0

for met, nu in rxn.metabolites.items():

s += float(nu) * float(lnC_vals[met.id])

delG[rxn.id] = float(delG0_dict[rxn.id]) + RT * s

return delG

\# ============================================================

\# MODEL BUILD

\# ============================================================

def build_and_solve(model, delG0, prot_cost, params):

Rg = params["R_kJ_per_mol_K"]

T = params["T_K"]

RT = Rg * T

x_lb = np.log(params["Cmin"])

x_ub = np.log(params["Cmax"])

BIG_M = params["BIG_M"]

m = gp.Model("ecOptMDFPathway")

m.setParam("OutputFlag", params["gurobi_OutputFlag"])

m.setParam("DualReductions", params["gurobi_DualReductions"])

\# --------------------------------------------------------

\# VARIABLES

\# --------------------------------------------------------

v = {}

y = {}

for rxn in model.reactions:

ub = max(0.0, float(rxn.upper_bound))

v[rxn.id] = m.addVar(lb=0, ub=ub, name=f"v_{rxn.id}")

y[rxn.id] = m.addVar(vtype=GRB.BINARY, name=f"y_{rxn.id}")

x = {met.id: m.addVar(lb=x_lb, ub=x_ub, name=f"x_{met.id}") for met in model.metabolites}

f = {rxn.id: m.addVar(lb=-GRB.INFINITY, name=f"f_{rxn.id}") for rxn in model.reactions}

B = m.addVar(lb=0, name="B")

m.update()

\# --------------------------------------------------------

\# MASS BALANCE

\# --------------------------------------------------------

for met in model.metabolites:

m.addConstr(

gp.quicksum(float(rxn.metabolites[met]) * v[rxn.id] for rxn in met.reactions) == 0,

name=f"mass_{met.id}",

)

\# --------------------------------------------------------

\# THERMODYNAMICS

\# --------------------------------------------------------

for rxn in model.reactions:

dG0 = float(delG0[rxn.id])

m.addConstr(

f[rxn.id] == -(dG0 + RT * gp.quicksum(float(coeff) * x[met.id] for met, coeff in rxn.metabolites.items())),

name=f"thermo_{rxn.id}",

)

\# --------------------------------------------------------

\# FLUX-ACTIVITY COUPLING

\# --------------------------------------------------------

for rxn in model.reactions:

ub = max(0.0, float(rxn.upper_bound))

m.addConstr(v[rxn.id] <= ub * y[rxn.id], name=f"flux_ub_{rxn.id}")

m.addConstr(v[rxn.id] >= params["min_flux"] * y[rxn.id], name=f"flux_lb_{rxn.id}")

\# --------------------------------------------------------

\# MDF COUPLING

\# --------------------------------------------------------

for rxn in model.reactions:

m.addConstr(f[rxn.id] >= B - BIG_M * (1 - y[rxn.id]), name=f"mdf_{rxn.id}")

\# --------------------------------------------------------

\# PROTEOME CONSTRAINT

\# --------------------------------------------------------

if prot_cost:

prot_expr = gp.quicksum(float(cost) * v[rid] for rid, cost in prot_cost.items()) / params["PROTEIN_COST_DIVISOR"]

m.addConstr(prot_expr <= params["PROTEOME_UB_g_per_gDW"], name="protein_capacity")

\# --------------------------------------------------------

\# MINIMUM BIOMASS

\# --------------------------------------------------------

biomass_id = params["BIOMASS_RXN"]

if params["min_biomass"] > 0:

m.addConstr(v[biomass_id] >= params["min_biomass"], name="min_biomass")

\# --------------------------------------------------------

\# OBJECTIVE

\# --------------------------------------------------------

total_flux = gp.quicksum(v[rxn.id] for rxn in model.reactions)

m.setObjective(v[biomass_id] + params["lambda_B"] * B - params["lambda_F"] * total_flux, GRB.MAXIMIZE)

m.optimize()

return m, v, x, B, RT

\# ============================================================

\# LOAD MODEL

\# ============================================================

model = cobra.io.read_sbml_model(PARAMS["MODEL_SBML"])

if PARAMS["BIOMASS_RXN"] not in [r.id for r in model.reactions]:

raise ValueError("Biomass reaction not found.")

\# ============================================================

\# LOAD ΔG°

\# ============================================================

delG_df = pd.read_excel(PARAMS["DELG_XLSX"])

delG0 = pd.Series(delG_df["delG0"].values, index=delG_df["Reaction"]).to_dict()

\# ============================================================

\# LOAD PROTEIN COST

\# ============================================================

prot_raw = pd.read_excel(PARAMS["PROTEIN_XLSX"], usecols=[0, 1], names=["Reactions", "Protein_cost"])

model_rxns = set(r.id for r in model.reactions)

prot_cost = {}

for _, row in prot_raw.iterrows():

rid = str(row["Reactions"])

if rid in model_rxns and pd.notna(row["Protein_cost"]):

prot_cost[rid] = float(row["Protein_cost"])

\# ============================================================

\# SOLVE

\# ============================================================

m, v, x, B, RT = build_and_solve(model, delG0, prot_cost, PARAMS)

if m.status != GRB.OPTIMAL:

raise RuntimeError("Optimization did not converge.")

\# ============================================================

\# EXTRACT SOLUTION

\# ============================================================

fluxes = {rid: var.X for rid, var in v.items()}

lnC = {mid: var.X for mid, var in x.items()}

C = {mid: np.exp(val) for mid, val in lnC.items()}

delG_model = compute_delG_from_solution(model, delG0, lnC, RT)

\# ============================================================

\# DATAFRAMES

\# ============================================================

flux_df = pd.DataFrame.from_dict(fluxes, orient="index", columns=["flux"]).reset_index()

flux_df.columns = ["Reaction", "flux"]

conc_df = pd.DataFrame.from_dict(C, orient="index", columns=["concentration"]).reset_index()

conc_df.columns = ["Metabolite", "concentration"]

thermo_df = pd.DataFrame({

"Reaction": [r.id for r in model.reactions],

"delG0_final": [delG0[r.id] for r in model.reactions],

"delG_model": [delG_model[r.id] for r in model.reactions],

})

\# ============================================================

\# PROTEIN USAGE

\# ============================================================

protein_rows = []

p_total = 0.0

for rid, cost in prot_cost.items():

pr = cost * fluxes.get(rid, 0) / PARAMS["PROTEIN_COST_DIVISOR"]

protein_rows.append({

"Reaction": rid,

"Protein_Required (g/gDW)": pr

})

p_total += pr

protein_df = pd.DataFrame(protein_rows)

\# ============================================================

\# WRITE EXCEL

\# ============================================================

with pd.ExcelWriter(PARAMS["OUTPUT_XLSX"], engine="openpyxl") as writer:

flux_df.to_excel(writer, sheet_name="flux", index=False)

conc_df.to_excel(writer, sheet_name="concentration", index=False)

thermo_df.to_excel(writer, sheet_name="thermodynamics", index=False)

protein_df.to_excel(writer, sheet_name="protein_requirements", index=False)

print("Biomass growth rate (/hr) =", v[PARAMS["BIOMASS_RXN"]].X)

print(f"Total protein requirement (g/gDW): {p_total}")

print("Minimum thermodynamic driving force (Kj/mol) =", B.X)

**ChatGEM Evaluation Report by Claude Sonnet 4.6: RAG vs. Without RAG**

Evaluated by Claude Sonnet 4.6

Overall Performance Score (OPS) = 0.7 x Coding Accuracy + 0.3 x Code Completeness

**1. Scoring Summary**

| **Task** | **Complexity** | **Condition** | **Accuracy (1-5)** | **Completeness (1-5)** | **OPS** |
| --- | --- | --- | --- | --- | --- |
| Task 1: FBA and Knockout | Low | Without RAG | 3.00 | 4.00 | 3.40 |
|  |  | With RAG | 4.00 | 5.00 | 4.30 |
| Task 2: OptKnock Strain Design | High | Without RAG | 1.00 | 2.00 | 1.30 |
|  |  | With RAG | 4.00 | 4.00 | 4.00 |
| Task 3: ecOptMDFPathway | Complex | Without RAG | 2.00 | 1.50 | 1.85 |
|  |  | With RAG | 4.00 | 5.00 | 4.30 |

**Scoring Rubric**

Coding Accuracy (1-5): 5 = runs correctly with expected output and follows best practices; 4 = runs correctly but with minor code quality or convention issues; 3 = runs but produces incorrect output; 2 = partially executes or uses wrong library; 1 = does not run.

Code Completeness (1-5): 5 = all required steps present; 4 = minor step or output omission; 3 = key step missing; 2 = major gaps; 1 = skeleton only.

**2. Findings by Task**

**Task 1: FBA and Knockout**

**Without RAG:** The code is structurally sound and correctly uses knock_out() on each target reaction. Biomass is correctly referenced via objective_value and succinate via reaction flux. However, the code retrieves the biomass reaction by hardcoded ID "BIOMASS" rather than the correct iJN1463 identifier "BIOMASS_KT2440_WT3", which would cause a KeyError at runtime. Succinate is retrieved via reaction flux rather than sol.fluxes["EX_succ_e"], which would also fail before the knockout. These two naming errors mean the code will raise an exception before completing the task, despite correct structural intent. Completeness is 4 because all protocol steps (baseline FBA, knockouts, post-knockout FBA, reporting) are represented.

**With RAG:** Verified functional. Uses exact EX_succ_e ID and knock_out() correctly. However, the model is modified in-place without a context manager, meaning knockouts are permanent on the model object. This is a minor code quality issue that does not affect correctness for this isolated task but would cause errors in a pipeline context.

**Task 2: OptKnock Strain Design**

**Without RAG:** The code imports from a fabricated strain_design package with nonexistent submodules (strain_design.model.CobraModel, strain_design.export.export_flux_comparison, strain_design.simulation.set_medium_exchanges). These do not exist in the actual straindesign library and the code will fail immediately on import. The glucose upper bound is incorrectly set to 0.0, which would block all glucose uptake and prevent any biomass growth. The OptKnock setup omits the sd.SDModule / sd.compute_strain_designs API and instead invents a bespoke ok.run() interface. These are fundamental errors rendering the code non-functional and non-compliant with the specified toolbox. Completeness is 2 because the logical structure covers all steps but the implementation is built on a fictional API.

**With RAG:** Verified functional. Uses the correct StrainDesign toolbox with proper sd.OPTKNOCK formulation. Minor issues: the exclusion list logic appends rxn_id to candidate_ko before evaluating the skip flag, meaning excluded reactions are still recorded in the candidate list even when the solution is skipped. Additionally, the flux comparison sheet includes a delta column, which is correctly computed and present.

**Task 3: ecOptMDFPathway**

**Without RAG:** The overall MILP structure is substantially correct: stoichiometric mass balance, thermodynamic dG calculation, flux-activity coupling via Big-M binary variables, proteome constraint, minimum biomass, and the weighted objective are all formulated and present. The use of abs(v[rxn]) in both the proteome constraint and objective is nonlinear and will cause Gurobi to reject the model unless the model is declared irreversible throughout. For the irreversible iJN1463_irrev.xml model this is partially justified, though the code does not enforce it explicitly. The code executes the full pipeline and exports all four required sheets. These issues are meaningful but do not prevent the code from attempting to solve—they reduce reliability rather than causing an immediate crash, so accuracy is 2.

**With RAG:** Verified functional. Clean Gurobi MILP with correct mass balance, flux-activity coupling, proteome constraint, and objective. Minor issue: the thermodynamic driving force variable f is defined as the negation of the standard dG expression, which is internally consistent but deviates from conventional sign convention, reducing interpretability. All output sheets are exported correctly.

**3. Overall Comparison**

| **Metric** | **Without RAG** | **With RAG** |
| --- | --- | --- |
| Mean Accuracy | 2.00 | 4.00 |
| Mean Completeness | 2.5 | 4.67 |
| Mean OPS | 2.18 | 4.20 |
