## Supplementary figures and images for "ChatGEM: An Agentic Architecture Enabling Interactive Simulation of Genome-Scale Metabolic Models" (from another version of this preprint: version 1, 2026-07-21)

### To evaluate whether model choice affects code generation performance, we compared the default o4-mini, a lightweight reasoning model, against GPT-4.1,

#### Slide 1
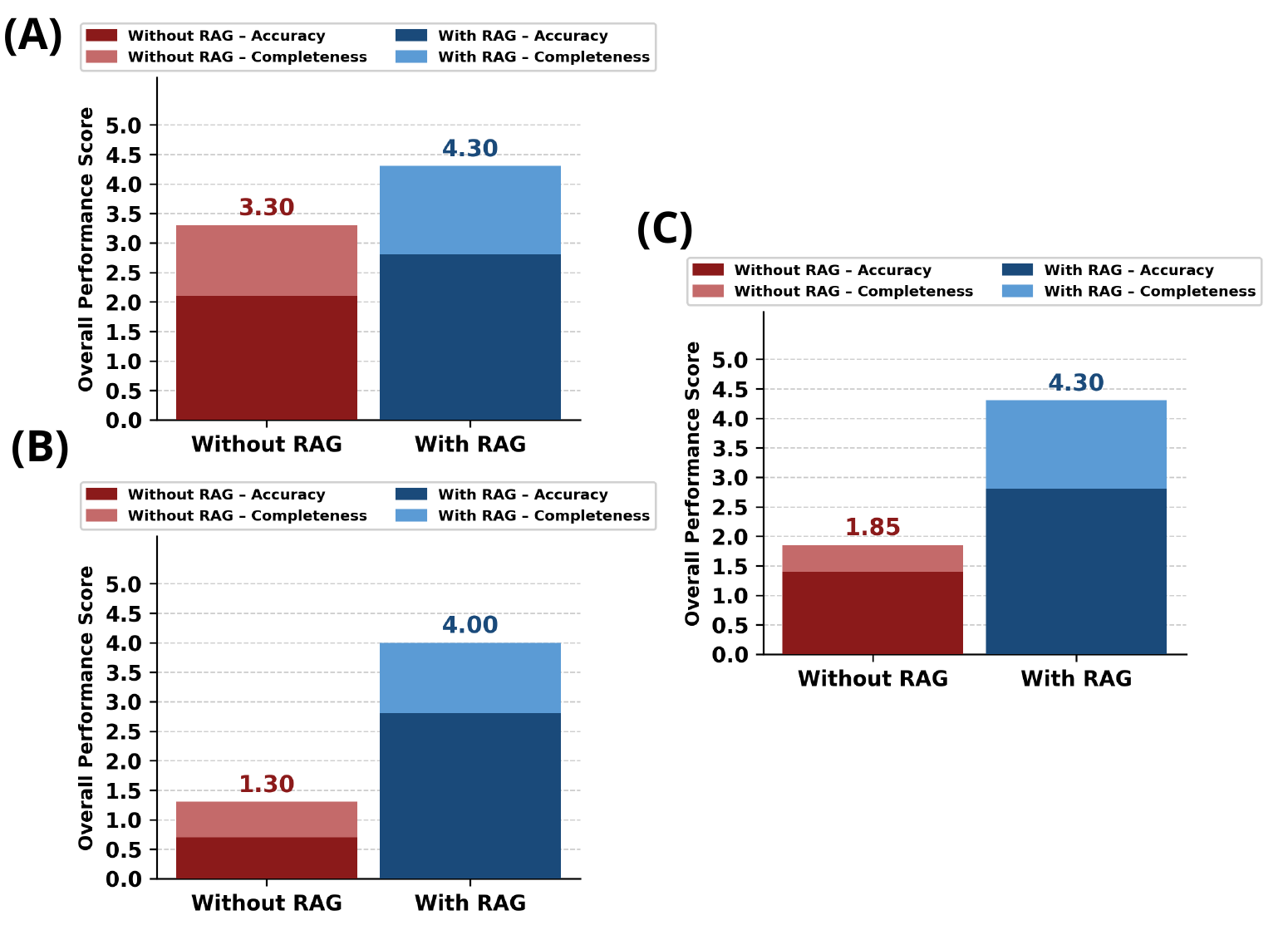

(A)
(C)
(B)
